# Fragmentation modes and the evolution of life cycles

**DOI:** 10.1101/120097

**Authors:** Yuriy Pichugin, Jorge Peña, Paul B. Rainey, Arne Traulsen

**Author notes:** Current Address: Institute for Advanced Study in Toulouse, 21 allée de Brienne, 31015 Toulouse Cedex 6, France (Jorge Peña).

## Abstract

Reproduction is a defining feature of living systems. To reproduce, aggregates of biological units (e.g., multicellular organisms or colonial bacteria) must fragment into smaller parts. Fragmentation modes in nature range from binary fission in bacteria to collective-level fragmentation and the production of unicellular propagules in multicellular organisms. Despite this apparent ubiquity, the adaptive significance of fragmentation modes has received little attention. Here, we develop a model in which groups arise from the division of single cells that do not separate but stay together until the moment of group fragmentation. We allow for all possible fragmentation patterns and calculate the population growth rate of each associated life cycle. Fragmentation modes that maximise growth rate comprise a restrictive set of patterns that include production of unicellular propagules and division into two similar size groups. Life cycles marked by single-cell bottlenecks maximise population growth rate under a wide range of conditions. This surprising result offers a new evolutionary explanation for the widespread occurrence of this mode of reproduction. All in all, our model provides a framework for exploring the adaptive significance of fragmentation modes and their associated life cycles.

**Author Summary:** Mode of reproduction is a defining trait of all organisms, including colonial bacteria and multicellular organisms. To produce offspring, aggregates must fragment by splitting into two or more groups. The particular way that a given group fragments defines the life cycle of the organism. For instance, insect colonies can reproduce by splitting or by producing individuals that found new colonies. Similarly, some colonial bacteria propagate by fission or by releasing single cells, while others split in highly sophisticated ways; in multicellular organisms reproduction typically proceeds via a single cell bottleneck phase. The space of possibilities for fragmentation is so vast that an exhaustive analysis seems daunting. Focusing on fragmentation modes of a simple kind we parametrise all possible modes of group fragmentation and identify those modes leading to the fastest population growth rate. Two kinds of life cycle dominate: one involving division into two equal size groups, and the other involving production of a unicellular propagule. The prevalence of these life cycles in nature is consistent with our null model and suggests that benefits accruing from population growth rate alone may have shaped the evolution of fragmentation mode.

## Introduction

A requirement for evolution – and a defining feature of life – is reproduction [1, 2, 3, 4]. Perhaps the simplest mode of reproduction is binary fission in unicellular bacteria, whereby a single cell divides and produces two offspring cells. In more complex organisms, such as colonial bacteria, reproduction involves fragmentation of a group of cells into smaller groups. Bacterial species demonstrate a wide range of fragmentation modes, differing both in the size at which the parental group fragments and the number and sizes of offspring groups [5]. For example, in the bacterium *Neisseria*, a diplococcus, two daughter cells remain attached forming a two-celled group that separates into two groups of two cells only after a further round of cell division [6]. *Staphylococcus aureus*, another coccoid bacterium, divides in three planes at right angles to one another to produce grape-like clusters of about 20 cells from which single cells separate to form new clusters [7]. Magnetotactic prokaryotes form spherical clusters of about 20 cells, which divide by splitting into two equally sized clusters [8].

These are just a few examples of a large number of diverse fragmentation modes, but why should there be such a wide range of life cycles? Do fragmentation modes have adaptive significance or are they simply the unintended consequences of particular cellular processes underpinning cell division? If adaptive, what selective forces shape their evolution? Can different life cycles simply provide different opportunities to maximise population growth rate?

A starting point to answer these questions is to consider benefits and costs of group living in cell collectives. Benefits may arise for various reasons. Cells within groups may be better able to withstand environmental stress [9], escape predation [10, 11], or occupy new niches [12, 13]. Also, via density-dependent gene regulation, cells within groups may gain more of a limiting resource than they would if alone [14, 15]. On the other hand, cells within groups experience increased competition and must also contend with the build up of potentially toxic waste metabolites [16, 17]. Thus, it is reasonable to expect an optimal relationship between group size and fragmentation mode that is environment and organism dependent [18, 19, 20, 21].

Here we formulate and study a matrix population model [22] that considers all possible modes of group fragmentation. By determining the relationship between life cycle and population growth rate, we show that there is, overall, a narrow class of optimal modes of fragmentation. When the process of fragmentation does not involve costs, optimal fragmentation modes are characterised by a deterministic schedule and binary splitting, whereby groups fragment into exactly two offspring groups. Contrastingly, when a cost is associated with fragmentation, it can be optimal for a group to fragment into multiple propagules.

Our results show that the range of life cycles observed in simple microbial populations are likely shaped by selection for intrinsic growth rate advantages inherent to different modes of group fragmentation. While we do not consider complex life cycles, our results can contribute to understanding the emergence of life cycles underpinning the evolution of multicellular life.

## Methods

### Group formation and fragmentation

We consider a population in which a single type of cell (or unit or individual) can form groups (or complexes or aggregates) of increasing size by cells staying together after reproduction [18]. We assume that the size of any group is smaller than *n*, and denote groups of size *i* by *X*_*i*_ (see the list of used variables in Table 1). Groups die at rate *d*_*i*_ and cells within groups divide at rate *b*_*i*_; hence groups grow at rate *ib*_*i*_. The vectors of birth rates **b** = (*b*_1_,…, *b*_*n*–1_) and of death rates d = (*d*_1_,…, *b*_*n*–1_) make the costs and benefits associated to the size of the groups explicit, thus defining the “fitness landscape” of our model.

**Table 1:** List of variables

|  |  |
| --- | --- |
| $X_i$ | a group of size $i$ cells |
| $x_i$ | abundance of groups of size $i$ |
| $\mathbf{x}$ | vector of abundances $x_i$ |
| $b_i$ | birth rate of cells in a group of $i$ cells |
| $d_i$ | death rates of groups of $i$ cells |
| $n$ | maximal group size |
| $\zeta_i$ | the number of partitions of integer $i$ |
| $\pi_i(\kappa)$ | the number of parts equal to $i$ in the partition $\kappa$ |
| $\mathbf{A}$ | projection matrix |
| $\lambda_1$ | population growth rate |
| $M$ | maximal benefit under monotonic fitness landscapes |
| $\alpha$ | degree of complementarity under monotonic fitness landscapes |

Groups produce new complexes by fragmenting (or splitting), i.e., by dividing into smaller groups. We further assume that fragmentation is triggered by growth of individual cells within a given group. Consider a group of size *i* growing into a group of size *i* + 1. Such a group can either stay together or fragment. If it fragments, it can do so in one of several ways. For example, a group of size 4 can give rise to the following five “fragmentation patterns”: 4 (the group does not split, but stays together), 3+1 (the group splits into two offspring groups: one of size 3, and one of size 1), 2+2 (the group splits into two groups of size 2), 2+1+1 (the group splits into one group of size 2 and two groups of size 1), and 1+1+1+1 (the group splits into four independent cells). Mathematically, such fragmentation patterns correspond to the five partitions of 4 (a partition of a positive integer *i* is a way of writing *i* as a sum of positive integers without regard to order; the summands are called parts [23]). We use the notation *κ* ⊦ *ℓ* to indicate that *κ* is a partition of *ℓ*, for example 2 + 2 ⊦ 4. The number of partitions of *ℓ* is given by *ζ_ℓ_*, e.g., there are *ζ*_4_ = 5 partitions of 4.

We consider an exhaustive set of fragmentation modes (or “fragmentation strategies”) implementing all possible ways groups of maximum size *n* can grow and fragment into smaller groups, including both pure and mixed modes (Fig. 1). A pure fragmentation mode is characterised by a single partition *κ* ⊦ *ℓ*, i.e., groups of size *i* < *ℓ* grow up to size *ℓ* and then fragment according to partition *κ* ⊦ *ℓ*. The partition *κ* can then be used to refer to the associated pure strategy. The total number of pure fragmentation strategies is 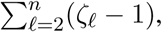 which grows quickly with *n*: There are 128 pure fragmentation modes for *n* = 10, but 1,295,920 for *n* = 50. A mixed fragmentation mode is given by a probability distribution over the set of pure fragmentation modes. The relationship between pure and mixed fragmentation modes is hence similar to the one between pure strategies and mixed strategies in evolutionary game theory [24]. One of our main results is that mixed fragmentation modes are always dominated by pure fragmentation modes. Hence, we focus our exposition on pure fragmentation modes, and leave the details of how to specify mixed fragmentation modes to Appendix A.

**Figure 1:**
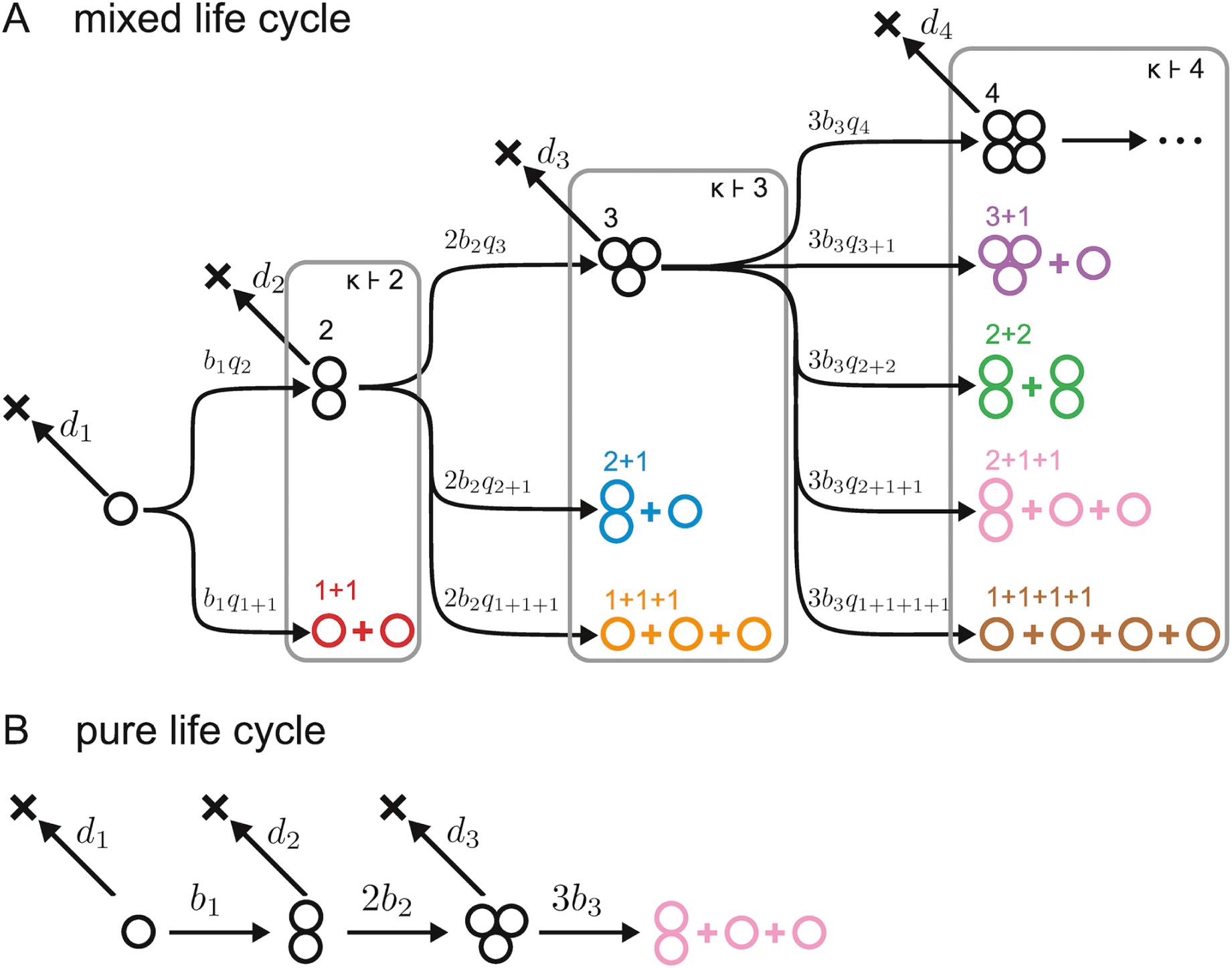
**Life cycles and fragmentation modes. A** Cells within groups of size *i* divide at rate *b*_*i*_, hence groups grow at rate *ib*_*i*_; groups die at rate *d*_*i*_. The sequences *b*_*i*_ and *d*_*i*_ define the fitness landscape of the model. We consider an exhaustive set of possible fragmentation modes, comprising both pure and mixed life cycles. In general, when growing from size *i* to size *i* + 1, groups stay together with probability *q*_*i*_, or fragment according to fragmentation pattern *κ* with probability *q*_*κ*_. Each fragmentation pattern (determining the number and size of offspring groups) can be identified with a partition of *i* + 1, i.e., a way of writing *i* + 1 as a sum of positive integers, that we denote by *κ* ⊦ *i* + 1. **B** Pure fragmentation modes are strategies with degenerate probability distributions over the set of partitions (so that *q*_*κ*_ = 1 for exactly one fragmentation pattern, including staying together). Here we illustrate the pure fragmentation mode 2 + 1 + 1, for which *q*_2_ = *q*_3_ = *q*2+1+1 = 1, and *q*_*κ*_ = 0 for all other *κ*.

### Biological reactions and population dynamics

Together with the fitness landscape given by the vectors of birth rates b and death rates d, each fragmentation strategy specifies a set of biological reactions. Consider the pure mode *κ* ⊦ *ℓ*, whereby groups grow up to size *ℓ* and then split according to fragmentation pattern *κ*. A set of reactions

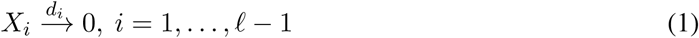

models the death of groups; an additional set of reactions

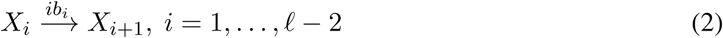

models the growth of groups (without splitting) up to size *ℓ* – 1. Finally, one reaction of the type

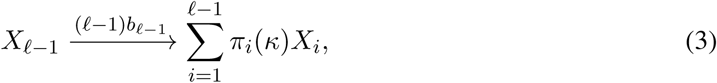

models the growth of the group from size *ℓ* – 1 to size *ℓ* and its immediate fragmentation in a way described by fragmentation pattern *κ* ⊦ *ℓ*, where parts equal to *i* appear a number *π*_*i*_(*κ*) of times. For instance, for the pure fragmentation mode 2 + 1 + 1 ⊦ 4, Eq. (3) becomes

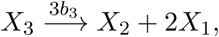

which stipulates that groups of size 3 grow to size 4 at rate 3*b*_3_ and split into one group of size 2 and two groups of size 1; here, *π*_1_(2 + 1 + 1) = 2, *π*_2_(2 + 1 + 1) = 1, *π*_3_(2 + 1 + 1) = 0.

The sets of reactions (1), (2), and (3) give rise to the system of differential equations

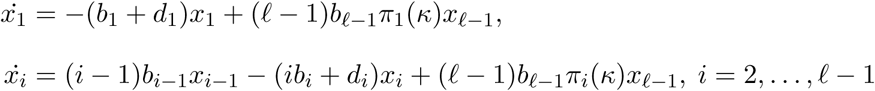

where *x*_*i*_ denotes the abundance of groups of size *i*. This is a linear system that can be represented in matrix form as

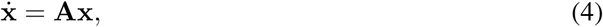

where x = (*x*_1_, *x*_2_,…, *x*_*ℓ*–1_) is the vector of abundances of the groups of different size and

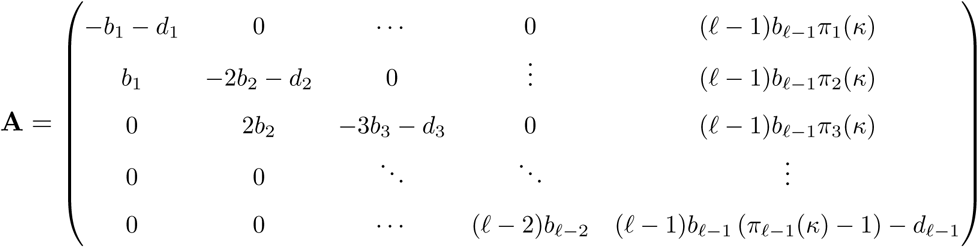

is the projection matrix determining the population dynamics.

### Population growth rate

For any fragmentation mode and any fitness landscape, the projection matrix **A** is “essentially non-negative” (or quasi-positive), i.e., all the elements outside the main diagonal are non-negative [25]. This implies that **A** has a real leading eigenvalue λ_1_ with associated non-negative left and right eigen-vectors v and w. In the long term, the solution of Eq. (4) converges to that of an exponentially growing population with a stable distribution, i.e.,

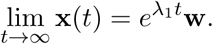

The leading eigenvalue λ_1_ hence gives the total population growth rate in the long term, and its associated right eigenvector **w** = (*w*_1_,…, *w*_*m*–1_) gives the stable distribution of group sizes so that, in the long term, the fraction *x*_*i*_ of complexes of size *i* in the population is proportional to *w*_*i*_.

### Dominance and optimality

For a given fitness landscape {**b**, **d**}, we can take the leading eigenvalue λ_1_(*κ*; **b**, **d**) as a measure of fitness of fragmentation mode *κ*, and consider the competition between two different fragmentation modes, *κ*_1_ and *κ*_2_. Indeed, under the assumption of no density limitation, the evolutionary dynamics are described by two uncoupled sets of differential equations of the form (4): one set for *κ*_1_ and one set for *κ*_2_. In the long term, *κ*_1_ is not outcompeted by *κ*_2_ if λ_1_(*κ*_1_; **b**, **d**) ≥ λ_1_(*κ*_2_; **b**, **d**); we then say that fragmentation mode *κ*_1_ dominates fragmentation mode *κ*_2_. We also say that strategy *κ*_*i*_ is optimal for given birth rates **b** and death rates **d** if it achieves the largest growth rate among all possible fragmentation modes.

### Two classes of fitness landscape: fecundity landscapes and survival landscapes

Fitness landscapes capture the many advantages or disadvantages associated with group living. These advantages may come either in the form of additional resources available to groups depending on their size or as an improved protection from external hazards. For our numerical examples, we consider two classes of fitness landscape, each representing only one of these factors. In the first class, that we call “fecundity landscapes”, group size affects only the birth rates of cells (while we impose *d*_*i*_ = 0 for all *i*). In the second class, that we call “survival landscapes”, group size affects only death rates (and we impose *b*_*i*_ = 1 for all *i*).

### Examples for *n* = 3

To fix ideas, consider all pure fragmentation modes with a maximum group size *n* = 3. These are 1+1 (“binary fission”, a partition of 2), 2+1 (“unicellular propagule”, a partition of 3), and 1+1+1 (“ternary fission” a partition of 3). The three associated projection matrices are given by

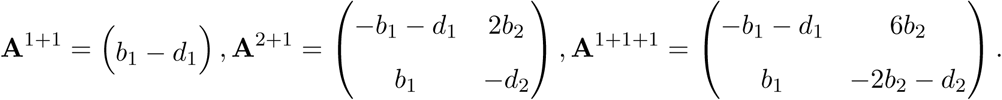

The three growth rates are

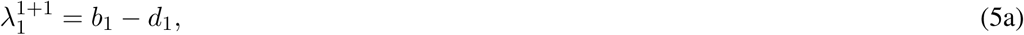

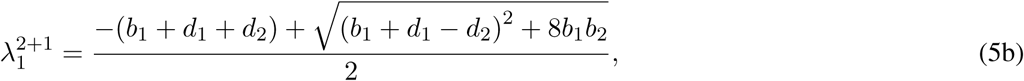

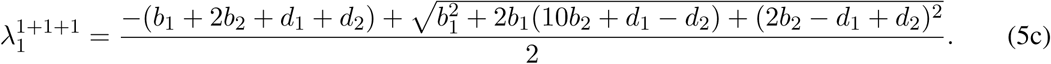

In the particular case of a fecundity landscape given by *b*_1_ = 1 and *b*_2_ = 15/8 (and *d*_1_ = *d*_2_ = 0), these growth rates reduce to 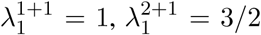 and 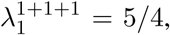 and we have 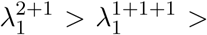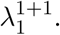 We then say that ternary and binary fission are dominated by the unicellular propagule strategy.

## Results

### Mixed fragmentation modes are dominated

Although for simplicity we focus our exposition on pure fragmentation strategies, we also consider mixed fragmentation strategies, i.e., probabilistic strategies mixing between different pure modes. A natural question to ask is whether a mixed fragmentation mode can achieve a larger growth rate than a pure mode. We find that the answer is no. For any fitness landscape and any maximum group size *n*, mixed fragmentation modes are dominated by a pure fragmentation mode (see Appendix B). Thus, the optimal fragmentation mode for any fitness landscape is pure.

As an example, consider fragmentation modes 1+1 and 2+1, and a mixed fragmentation mode mixing between these two so that with probability *q* splitting follows fragmentation pattern 2+1 and with probability 1 – *q* it follows fragmentation pattern 1+1. For any mixing probability *q* and any fitness landscape, the growth rate of the mixed fragmentation mode is given by

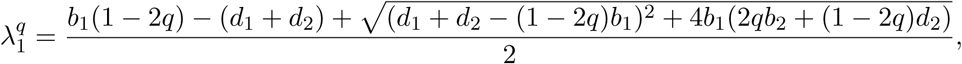

which can be shown to always lie between the growth rates of the pure fragmentation modes, i.e., either 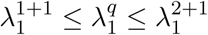 or 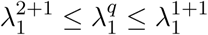 holds and the mixed fragmentation mode is dominated (see Appendix C).

To further illustrate our analytical findings, consider groups of maximum size *n* = 4 and a fecundity landscape given by b = (1, 2, 1.4). We randomly generated 10^7^ mixed fragmentation modes by drawing the probabilities for growth without splitting from an uniform distribution and letting the probabilities of splitting according to a given fragmentation pattern be proportional to exponential random variables with rate parameter equal to one. We then calculated the growth rate of these mixed strategies together with the growth rate of the seven pure fragmentation modes available for *n* = 4, i.e., 1+1, 2+1, 1+1+1, 3+1, 2+2, 2+1+1, and 1+1+1+1 (Fig. 2A). In line with our analysis, a pure fragmentation mode (namely 2+2, whereby groups grow up to size 4 and then immediately split into two bicellular groups) achieves a higher growth rate than the growth rate of any mixed fragmentation mode, and the highest growth rate overall.

**Figure 2:**
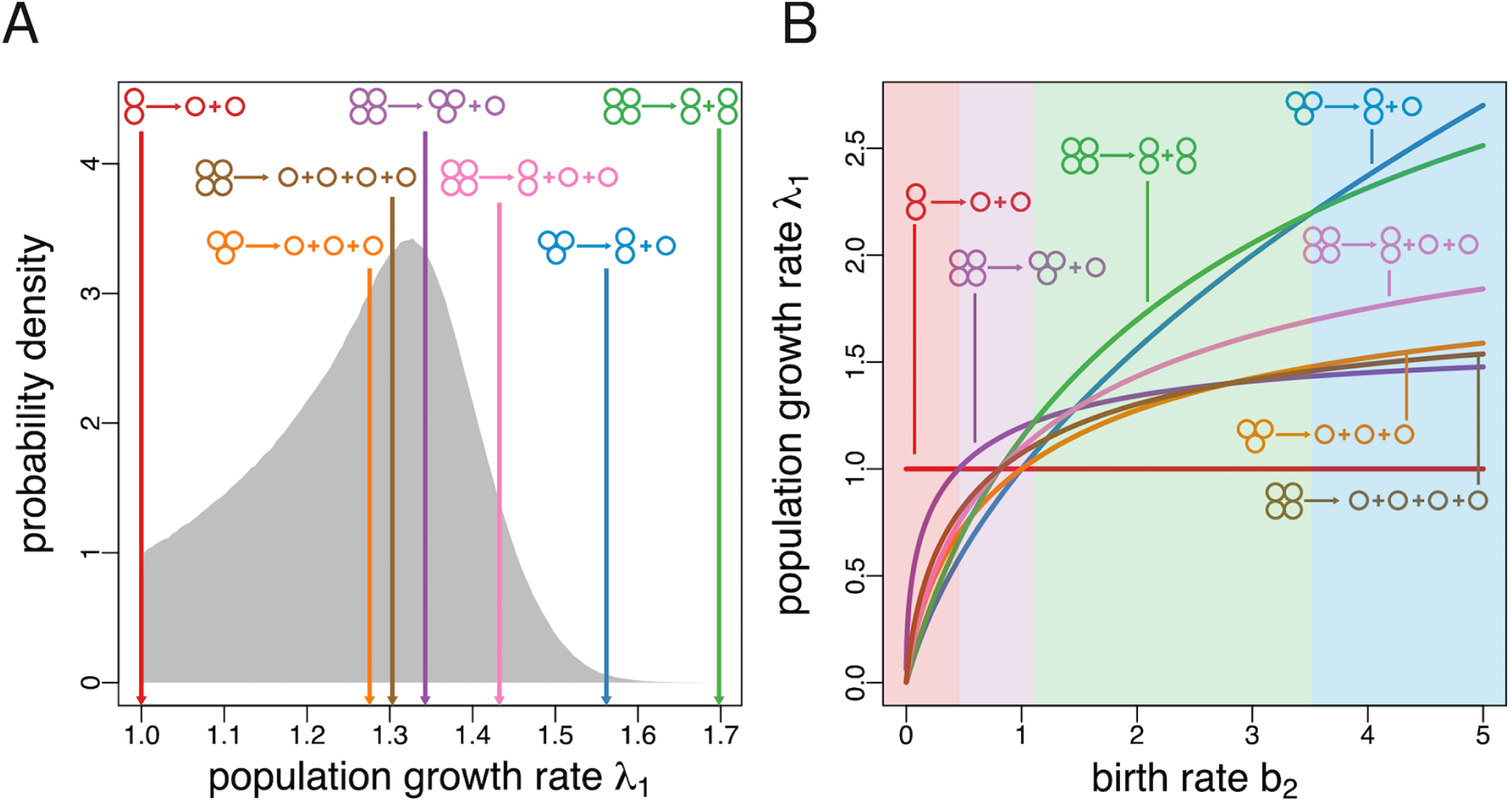
**The optimal fragmentation mode is pure and characterised by binary fragmentation. A** Mixed fragmentation strategies are dominated. Here we show the empirical probability distribution of the growth rate of mixed fragmentation modes for *n* = 4 (generated from a sample of 10^7^ randomly generated fragmentation modes) subject to the fitness landscape {**b**, **d**} = {(1, 2, 1.4), (0, 0, 0)}. The growth rates of all seven pure fragmentation modes for *n* = 4 are indicated by arrows. In this case, 2+2 achieves the maximal possible growth rate among all possible fragmentation modes. **B** Optimal fragmentation modes are characterised by binary splitting. Population growth rate (λ_1_) for all seven pure fragmentation modes for *n* = 4 subject to the fitness landscape {**b**, **d**} = {(1, *b*_2_, 1.4), (0, 0, 0)} as a function of the birth rate of groups of size 2, *b*_2_. Each of the four fragmentation modes characterised by binary fragmentation (1+1, 2+1, 2+2, and 3+1) can be optimal depending on the value of *b*_2_. Contrastingly, nonbinary fragmentation modes (1+1+1, 1+1+1+1, and 2+1+1) are never optimal.

### Optimal fragmentation modes are characterised by binary splitting

Having shown that mixed fragmentation modes are dominated, we now ask which pure modes might be optimal. We find that, within the set of pure modes, “binary” fragmentation modes (whereby groups split into exactly two offspring groups) dominate “nonbinary” fragmentation modes (whereby groups split into more than two offspring groups). To illustrate this result, consider the simplest case of *n* = 3 and the three modes 1+1, 2+1, and 1+1+1, out of which 1+1 and 2+1 are binary, and 1+1+1 is nonbinary. Comparing their growth rates (as given in Eq. (5)), we find that 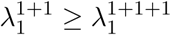 holds if *b*_1_ – *b*_2_ ≥ *d*_1_ – *d*_2_ and that 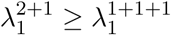 holds if *b*_1_ – *b*_2_ ≤ *d*_1_ – *d*_2_. Thus, for any fitness landscape, 1+1+1 is dominated by either 2+1 or by 1+1. More generally, we can show that for any nonbinary fragmentation mode, one can always find a binary fragmentation mode achieving a greater or equal growth rate under any maximum group size *n* and fitness landscape (see Appendices D and E). Taken together, our analytical results imply that the set of optimal fragmentation modes is countable and, even for large *n*, relatively small. Consider the proportion of pure fragmentation modes that can be optimal, which is defined by the ratio between the number of binary fragmentation modes and the total number of pure fragmentation modes. While this ratio is relatively high for small *n* (e.g., 2/3 ≈ 0.67 for *n* = 3 or 4/7 ≈ 0.57 for *n* = 4), it decreases sharply with increasing *n* (e.g., 25/128 ≈ 0.20 for *n* = 10 and 625/1295920 ≈ 0.00048 for *n* = 50).

Fig. 2B shows the growth rate of the seven pure modes for *n* = 4 for a fecundity landscape given by **b** = (1, *b*_2_, 1.4) as a function of *b*_2_. In line with our analysis, only binary fragmentation modes (1+1, 2+1, 2+2, and 3+1) can be optimal, while nonbinary fragmentation modes (1+1+1, 2+1+1, and 1+1+1+1) are dominated. Which particular binary mode is optimal depends on the particular value of the birth rate of groups of two cells. For small values (*b*_2_ ≲ 0.45), the fecundity of such groups is too low, and the optimal fragmentation mode is 1+1. For intermediary values (0.45 ≲ *b*_2_ ≲ 1.11), the reproduction efficiency of groups of three cells mitigates the inefficiency of cell pairs, and the mode 3+1 becomes optimal. For larger values (1.11 ≲ *b*_2_ ≲ 3.52), the optimal fragmentation mode is 2+2, where no single cells are produced. Finally, for very large values (*b*_2_ ≳ 3.52), the optimal fragmentation mode is 2+1; this ensures that one offspring group emerges at the most productive bicellular state.

More generally, which particular fragmentation mode within the class of binary splitting strategies is optimal depends on all birth rates and death rates characterising the fitness landscape. To further explore this issue, we identified the optimal fragmentation modes for general fecundity and survival landscapes for the simple case of *n* = 4 (Fig. 3; Appendix F). Since we can set *b*_1_ = 1 and min(**d**) = 0 without loss of generality (see Appendix D), we represent fitness landscapes as points in a two-dimensional parameter space with coordinates *b*_2_*/b*_1_ and *b*_3_*/b*_1_ for fecundity landscapes, and coordinates *d*_2_ – *d*_1_ and *d*_3_ – *d*_1_ for survival landscapes. The exact boundaries of the parameter regions where a given fragmentation mode is optimal are often nontrivial mathematical expressions. Nevertheless, we identify general patterns dictating which fragmentation mode will be optimal. Consider first the optimality map for fecundity landscapes (Fig. 3A). A sufficient condition for the unicellular life cycle 1+1 to be optimal is that the birth rate of single cells is larger than the birth rate of pairs and triplets of cells (*b*_1_ > *b*_2_ and *b*_1_ > *b*_3_). In this case, there is no apparent reason why a fragmentation mode different than 1+1 would be optimal. Perhaps less trivially, 1+1 can also be optimal in cases where single cells are less fertile than groups of three cells, i.e., even if *b*_1_ < *b*_3_ holds. This requires the birth rate *b*_2_ to be so small that the fecundity benefits accrued when reaching the size of three cells are not enough to compensate for the unavoidable penalty of passing through the less prolific state of two cells. Turning now to fragmentation mode 2+1, a necessary condition for this mode to be optimal is that pairs of cells have the largest birth rate, i.e., that *b*_2_ > *b*_1_ and *b*_2_ > *b*_3_ holds. Similarly, mode 3+1 can only be optimal if *b*_3_ > *b*_1_ and *b*_3_ > *b*_2_, so that groups of three have the largest birth rate. In these two cases, the optimal fragmentation mode (either 2+1 or 3+1) keeps one of the two offspring groups at the most productive size. Finally, for fragmentation mode 2+2 to be optimal, it is necessary that single cells have the lowest birth rate, i.e., that *b*_2_ > *b*_1_ and *b*_3_ > *b*_1_ holds. In this case, the fragmentation mode ensures that the life cycle of the organism never goes through the least productive unicellular phase. Under survival landscapes, fitness increases as death rates decrease. Taking this qualitative difference into account, the map of optimal fragmentation modes under survival landscapes (Fig. 3B) follows similar qualitative patterns as the one under fecundity landscapes.

**Figure 3:**
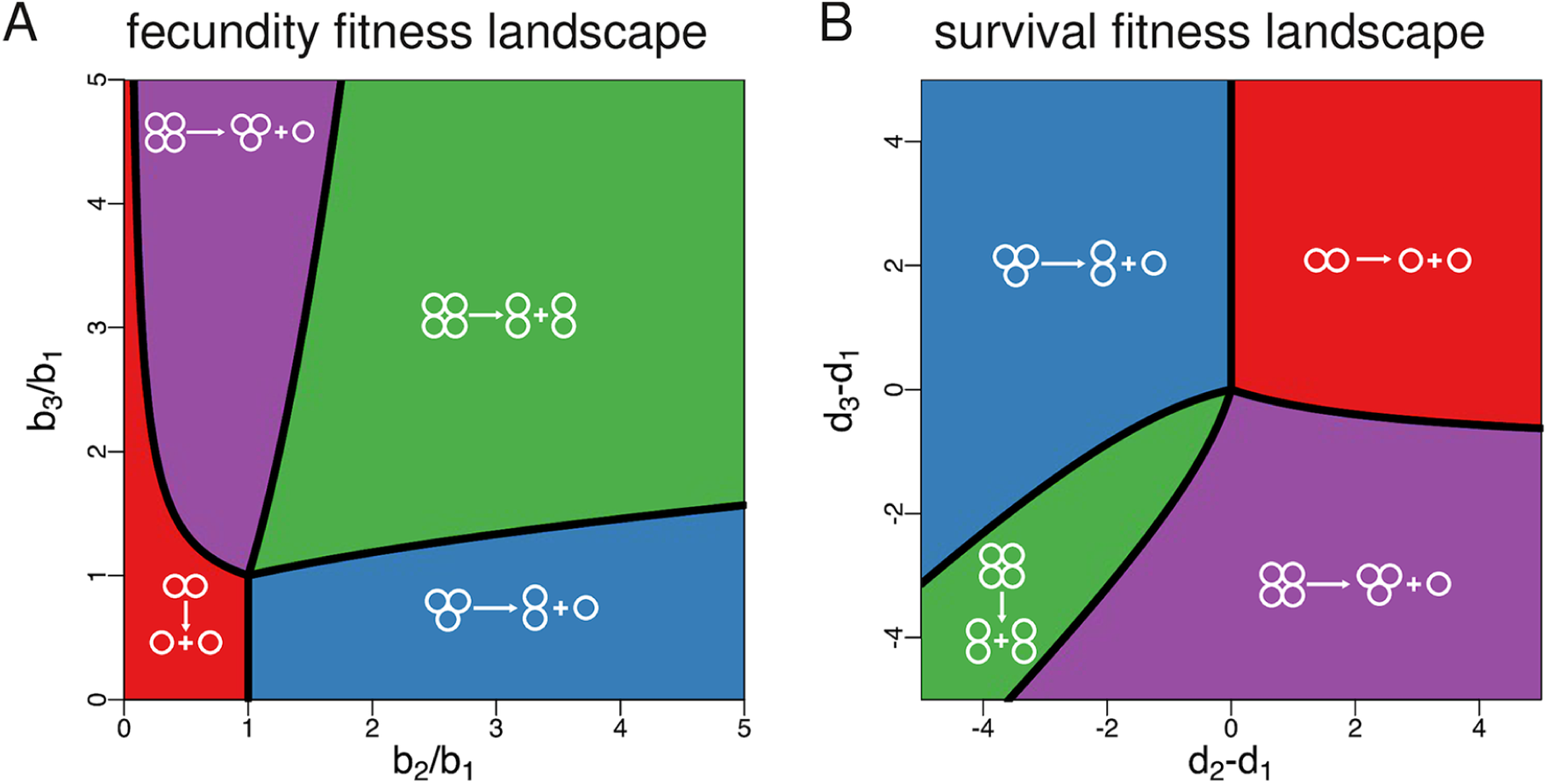
**Optimal fragmentation modes for fecundity and survival landscapes (costless fragmentation). A** Life cycles achieving the maximum population growth rate for *n* = 4 under fecundity landscapes (i.e., *d*_1_ = *d*_2_ = *d*_3_ = 0). In this scenario, fragmentation mode 2+2 is optimal for most fitness landscapes. **B** Life cycles achieving the maximum population growth rate for *n* = 4 under survival landscapes (i.e., *b*_1_ = *b*_2_ = *b*_3_ = 1). In this scenario, fragmentation modes emitting a unicellular propagule (1+1, 2+1, 3+1) are optimal for most parameter values. We use ratios of birth rates and differences between death rates as axes because one can consider *b*_1_ = 1 and min(*d*_1_, *d*_2_, *d*_3_) = 0 without loss of generality (see Appendix D). Shaded areas are obtained from the direct comparison of numerical solutions, lines are found analytically (see Appendix F).

### Costly fragmentation allows for optimal nonbinary fragmentation and multicellularity without group benefits

So far we have assumed that fragmentation is costless. However, fragmentation processes can be costly to the parental group undergoing division. This is particularly apparent in cases where some cells need to die in order for fragmentation of the group to take place. Examples in simple multicellular forms include *Volvox*, where somatic cells constituting the outer layer of the group die upon releasing the offspring colonies and are not passed to the next generation [26], the breaking of filaments in colonial cyanobacteria [27], and the fragmentation of “snowflake-like” clusters of the yeast *Saccharomyces cerevisiae* [28]. Fragmentation costs may also be less apparent. For instance, fragmentation may cost resources that would otherwise be available for the growth of cells within a group.

To investigate the effect of fragmentation costs on the set of optimal fragmentation modes, we consider two cases: proportional costs and fixed costs. For proportional costs, we assume that *π* – 1 cells die in the process of a group fragmenting into *π* parts. This case captures the fragmentation process of filamentous bacteria, where filament breakage entails the death of cells connecting the newly formed fragments [27]. For fixed costs, we assume that exactly one cell is lost upon each fragmentation event. This scenario is loosely inspired by yeast colonies with a tree-like structure, where cells can be connected with many other cells, so the death of a single cell may release more than two offspring colonies [28, 19]. Mathematically, both cases imply that fragmentation patterns are described by partitions of a number smaller than the size of the parent group (see Appendix G).

For both kinds of costly fragmentation, we can show that mixed fragmentation modes are still dominated by pure fragmentation modes (the proof given in Appendix B also holds in this case). Moreover, for proportional costs the optimal fragmentation mode is also characterised by binary fragmentation, as it is the case for costless fragmentation (see Appendix H). This makes intuitive sense, as the addition of a penalty for splitting into many fragments should further reinforce the optimality of binary splitting (whereby only one cell per fragmentation event is lost). In contrast, we find that under fragmentation with fixed costs the optimal fragmentation mode can involve nonbinary fragmentation, i.e., division into more than two offspring groups. This result can be readily illustrated for the case of *n* = 4 where the nonbinary mode 1+1+1 is optimal for a wide range of fitness landscapes (Fig. 4).

**Figure 4:**
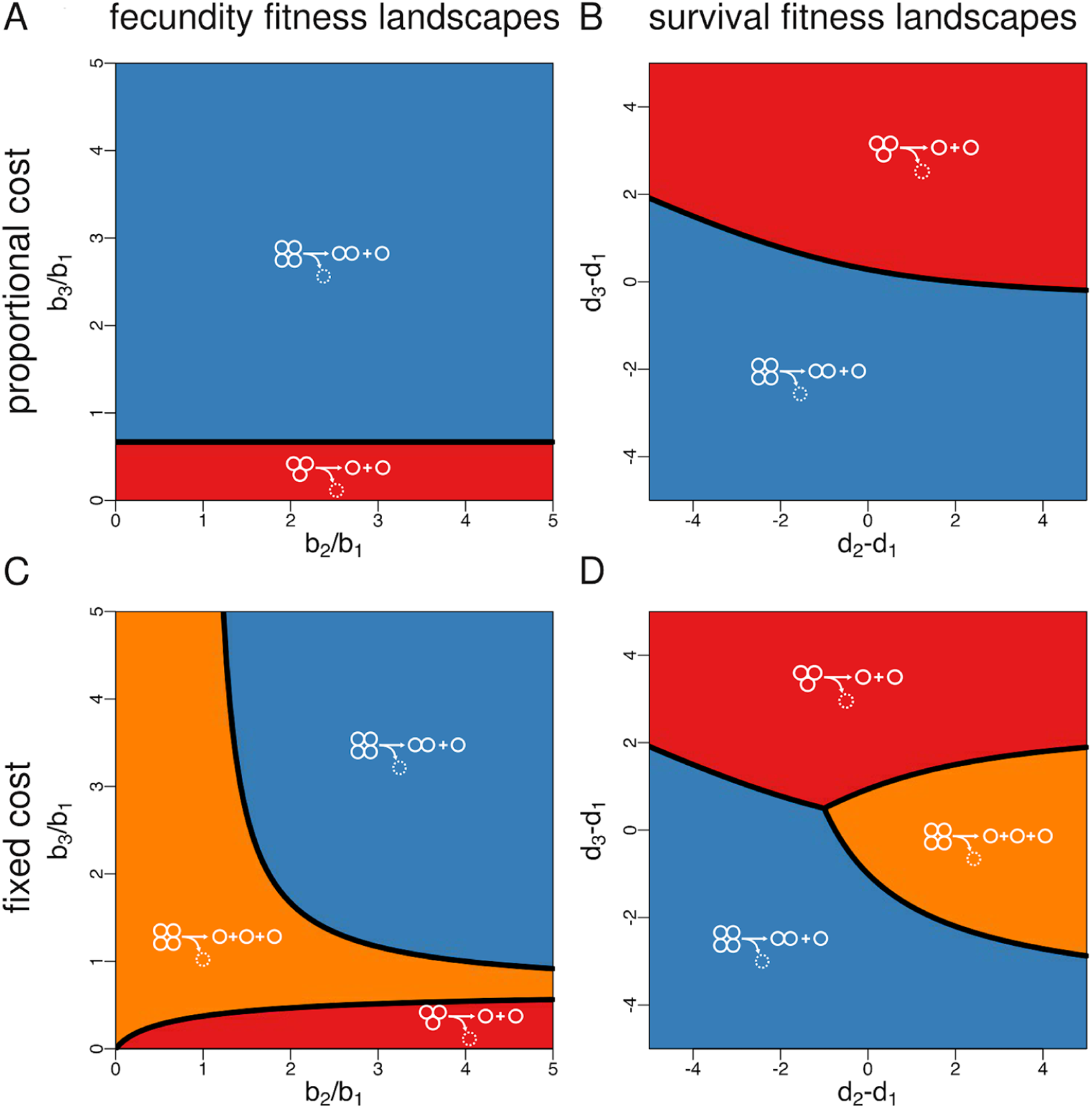
**Optimal fragmentation modes for fecundity and survival landscapes (costly fragmentation).** For proportional costs (panels **A** and **B**), splitting into *π* parts involves the loss of *π* – 1 cells. In this case, and for *n* = 4, only two pure modes are possible: 2+1 (whereby a 4-unit group splits into a pair of cells and a single cell and loses one cell) and 1+1 (whereby a group of three splits into two single cells and loses one cell). For fixed costs (panels **C** and **D**), splitting involves the loss of a single cell, no matter the kind of partition. In this case, and for *n* = 4, an additional mode is possible: 1+1+1 (whereby a 4-unit group splits into three single cells and loses one cell). This last nonbinary mode can be optimal under a wide range of parameters.

Another interesting feature of costly fragmentation (implemented via either proportional or fixed costs) is that fragmentation modes involving the emergence of large groups can be optimal even if being in a group does not grant any fecundity or survival advantage to cells. If fragmentation is costless, as we assumed before, fitness landscapes for which groups perform worse than unicells (that is, *b*_*i*_/*b*_1_ ≤ 1 for fecundity landscapes or *d*_*i*_ – *d*_1_ ≥ 0 for survival landscapes) lead to optimal fragmentation modes where splitting occurs at the minimum possible group size *i* = 2, so that no multicellular groups emerge in the population (cf. Fig. 3). In contrast, under costly fragmentation some of these fitness landscapes allow for the evolutionary optimality of fragmentation modes according to which groups split at the maximum size *n* = 4 (2+1 under proportional costs, and 1+1+1 under fixed costs), and hence for life cycles where multicellular phases are persistent. This seems paradoxical until one realises that by staying together as long as possible groups delay as much as possible the inevitable cell loss associated to a fragmentation event. Thus, even if groups are less fecund or die at a higher rate than independent cells, staying together might be adaptive if splitting apart is too costly.

### Synergistic interactions between cells promote the production of unicellular propagules, while diminishing returns promote equal binary fragmentation

Next, we focus on fitness landscapes for which either the birth rate of cells increases with group size (fecundity landscapes where larger groups are always more productive) or the death rate of groups decreases with group size (survival landscapes where larger groups always live longer). In this case, and for a maximum group size *n* = 4, the set of optimal modes is given by 2+2 and 3+1 if there are no fragmentation costs (Fig. 3), by 2+1 if fragmentation costs are proportional to the number of fragments (Fig. 3A-B), and by 2+1 and 1+1+1 if fragmentation involves a fixed cost of one cell (Fig. 4C-D).

To investigate larger maximum group sizes *n* in a simple but systematic way, we consider fecundity landscapes with birth rates given by *b*_*i*_ = 1 + *Mg*_*i*_ and survival landscapes with death rates given by *d*_*i*_ = *M*(1 – *g*_*i*_), where *g*_*i*_ = [(*i* – 1)/(*n* – 2)]^*α*^ [29] models the fecundity or survival benefits associated to group size *i* and *M* > 0 is the maximum benefit (Fig. 5). The parameter *α* is the degree of complementarity between cells; it measures how important the addition of another cell to the group is in producing the maximum possible benefit *M* [30]. For low degrees of complementarity (*α* < 1), the sequence *g*_*i*_ is strictly concave and each additional cell contributes less to the per capita benefit of group living [31] and groups of all sizes achieve the same functionality as *α* tends to zero. If *α* = 1, the sequence *g*_*i*_ is linear, and each additional cell contributes equally to the fecundity or survival of the group. Finally, for high degrees of complementarity (*α* > 1), the sequence *g*_*i*_ is strictly convex and each additional cell improves the performance of the group more than the previous cell did. In the limit of large *α*, the advantages of group living materialise only when complexes achieve the maximum size *n* – 1 [31].

**Figure 5:**
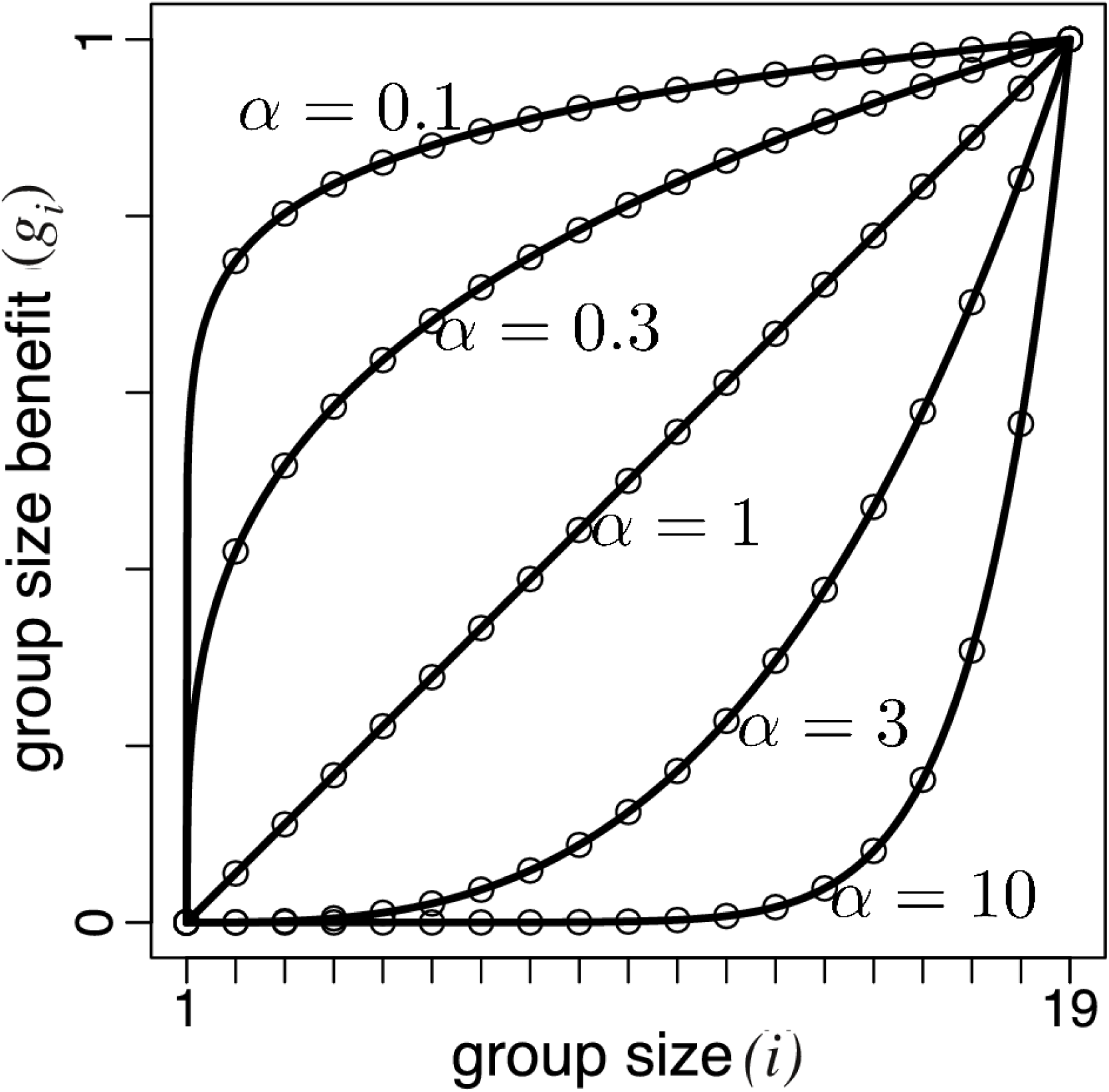
**Advantages of group living.** Group size benefit *g*_*i*_ = [(*i* – 1)/(*n* – 2)]^*α*^ as a function of group size for different values of the degree of complementarity *α*. If *α* < 1, *g*_*i*_ is concave; if *α* = 1, *g*_*i*_ is linear; if *α* > 1, *g*_*i*_ is convex.

We numerically calculate the optimal fragmentation modes for *n* = 20 (costless fragmentation) or *n* = 21 (costly fragmentation) and the fitness landscapes described above for parameter values taken from 0.01 ≤ *α* ≤ 100 and 0.02 ≤ *M* ≤ 50 (Figs. 6 and 7). In line with our general analytical results, optimal fragmentation modes are always characterised by binary splitting when fragmentation is costless or when it involves proportional costs, while nonbinary splitting can be optimal only if fragmentation involves a fixed cost. We also find that, for each value of *α* and *M*, and for both costless and costly fragmentation, the optimal fragmentation mode is one where fragmentation occurs at the largest possible size. This is expected since the benefit sequence is increasing in group size and thus groups of maximum size perform better, either by achieving the largest birth rate per unit (fecundity landscapes) or the lowest death rate (survival landscapes). Which particular fragmentation mode maximizes the growth rate depends nontrivially on whether fragmentation is costless or costly (and in the latter case also on how such costs are implemented), on the kind of group size benefits (fecundity or survival), on the maximum possible benefit *M*, and on the degree of complementarity *α*. Despite this apparent complexity, some general patterns can be identified.

**Figure 6:**
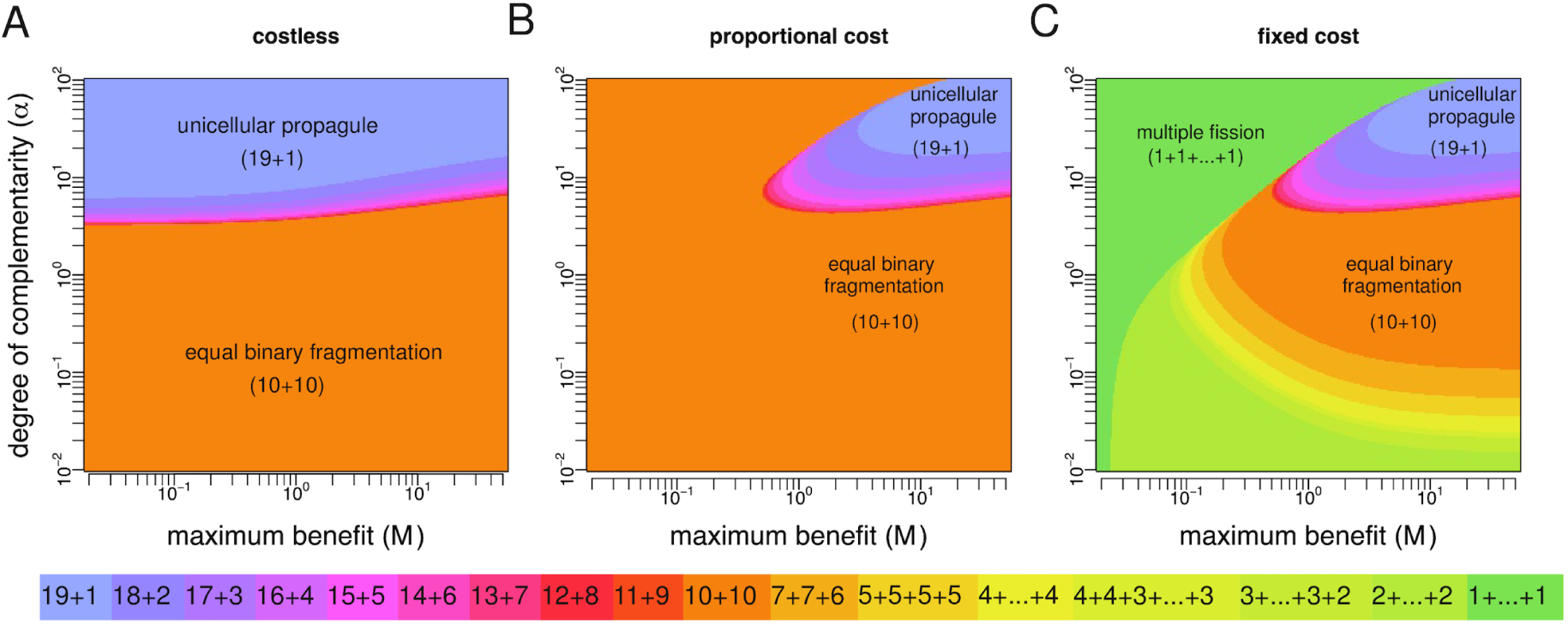
**Optimal life cycles under monotonic fecundity landscapes.** Birth rates are given by *b*_*i*_ = 1 + *Mg*_*i*_ where *g*_*i*_ = [(*i* – 1)/(*n* – 2)]^*α*^. **A** Costless fragmentation, *n* = 20. **B** Fragmentation with proportional costs, *n* = 21. **C** Fragmentation with fixed costs, *n* = 21. For costless fragmentation and fragmentation with proportional costs, only binary modes 19+1, 18+2,…, 10+10 are optimal. In these cases, diminishing returns (*α* < 1) make equal binary fragmentation (10+10) optimal. Also, optimality of the unicellular propagule strategy (19+1) requires increasing returns (*α* > 1). For fragmentation with fixed costs, nonbinary modes 7+7+6,…,1+…+1 can also be optimal.

**Figure 7:**
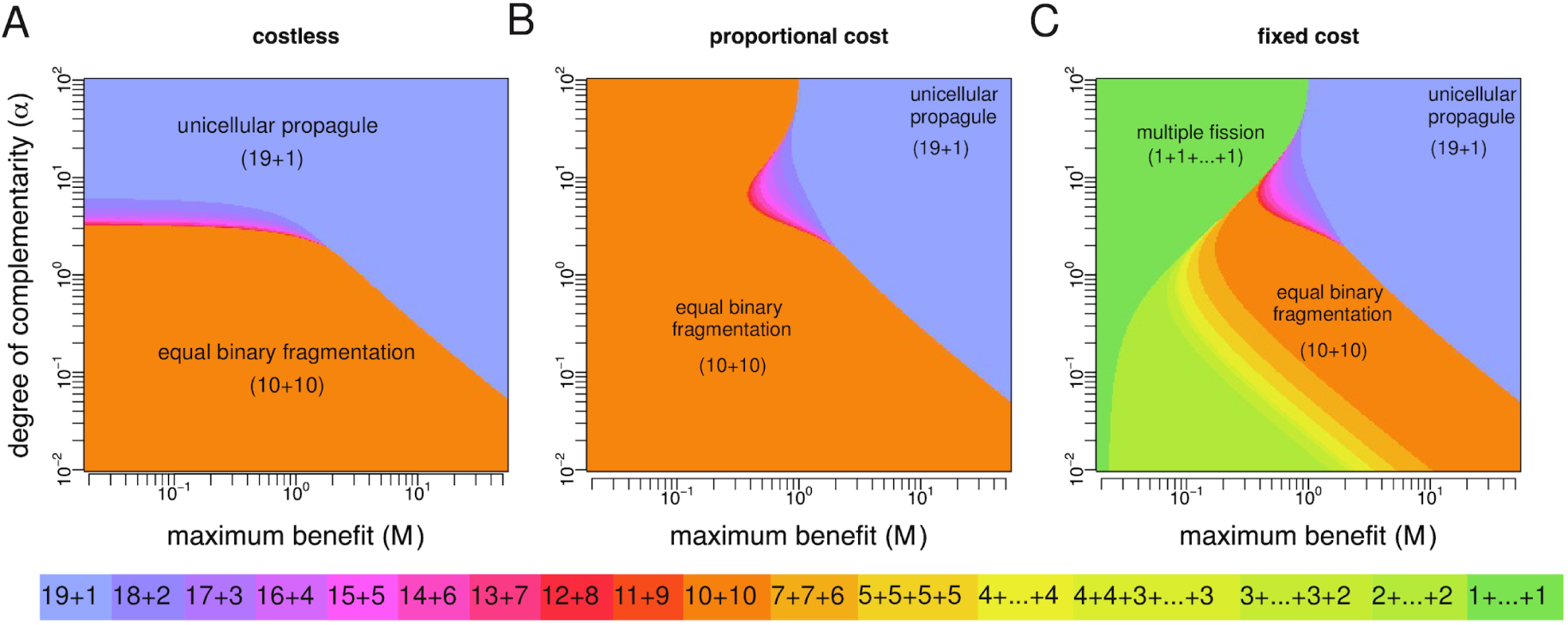
**Optimal life cycles under monotonic survival landscapes.** Death rates are given by *d*_*i*_ = *M*(1 – *g*_*i*_) where *g*_*i*_ = [(*i* – 1)/(*n* – 2)]^*α*^. **A** Costless fragmentation, *n* = 20. **B** Fragmentation with proportional costs, *n* = 21. **C** Fragmentation with fixed costs, *n* = 21. For costless fragmentation and fragmentation with proportional costs, only binary modes 19+1, 18+2,…, 10+10 are optimal. In these cases, diminishing returns to scale (*α* < 1) make equal binary fragmentation (10+10) optimal. Also, optimality of the unicellular propagule strategy (19+1) requires increasing returns to scale (*α* > 1). For fragmentation with fixed costs, nonbinary modes 7+7+6,…,1+…+1 can also be optimal.

Let us focus on the case of fecundity landscapes and first fasten attention on the scenario of costless fragmentation (Fig. 6A). A salient feature of this case is the prominence of two qualitatively different fragmentation modes: the “equal binary fragmentation” strategy 10+10 (whereby offspring groups have sizes as similar as possible) and the “unicellular propagule” strategy 19+1 (whereby offspring groups have sizes as different as possible). A sufficient condition for equal binary fragmentation to be optimal is that increase in size is characterised by diminishing returns. The intuition behind this result is that, if the degree of complementarity is small, then groups (complexes of size *i* ≥ 2) have similar performance, while independent cells (*i* = 1) are at a significant disadvantage. Therefore, the optimal strategy is to ensure that both offspring groups are as large as possible, and hence of the same size. However, equal binary fragmentation can be also optimal for synergistic interactions, provided that complementarity is not too high. In contrast, the unicellular propagule strategy is optimal only for relatively high degrees of complementarity. This is because when complementarity is high only the largest group can reap the benefits of group living; in this case, the optimal mode is to have at least one offspring of very large size. Compared to 19+1 and 10+10, other binary splitting strategies are optimal in smaller regions of the parameter space, and in all cases only for synergetic interactions between cells.

Consider now the effects of introducing fragmentation costs proportional to the number of fragments (Fig. 6B). Here, the region where the unicellular propagule strategy is optimal shrinks to the corner of the parameter space where benefits of group living and degree of complementarity are maximum, while the region of optimality for equal binary fragmentation expands. This makes intuitive sense. With fragmentation costs, the largest offspring group resulting from fragmenting according to the unicellular propagule strategy is of size 19, and hence always on the brink of fragmentation (once it grows to size 21) and incurring one cell loss. When group benefits are not high and synergistic enough, the unicellular propagule strategy is dominated by fragmentation modes (in particular, equal binary fragmentation) having smaller offspring for which the costs of fragmentation are not so immediate.

Finally, if costs of fragmentation are not proportional but fixed (Fig. 6C), then two classes of nonbinary splitting become optimal in regions of the parameter space where equal binary fragmentation was optimal under proportional costs: (i) “multiple fission” (1+1+…+1) which is in general favored for small maximum benefit and increasing returns, and (ii) “multiple groups” (modes 2+2+…+2, 3+3+3+3+3+3+2, 4+4+3+3+3+3, 4+4+4+4+4, 5+5+5+5, and 7+7+6) which are often optimal for diminishing returns.

Fig. 7 show the results for survival landscapes. The main difference in this case is that the unicellular propagule strategy can be the optimal strategy even when group living is characterised by diminishing returns. In general, fecundity benefits make equal binary fragmentation optimal under more demographic scenarios, while survival benefits make the unicellular propagule strategy optimal under more demographic scenarios.

## Discussion

Reproduction is such a fundamental feature of living systems that the idea that the mode of reproduction may be shaped by natural selection is easily overlooked. Here, we analysed a matrix population model that captures the demographic dynamics of complexes that grow by staying together and reproduce by fragmentation. The costs and benefits associated with group size ultimately determine whether or not a single cell fragments into two separate daughter cells upon cell division, or whether those daughter cells remain in close proximity, with fragmentation occurring only after subsequent rounds of division. We allowed for a vast and complete space of fragmentation strategies, including pure modes (specifying all possible combinations of size at fragmentation and fragmentation pattern) and mixed modes (specifying all probability distributions over the set of pure modes), and identified those modes achieving a maximum growth rate for given fecundity and survival size-dependent rates. Our research questions and methodology thus resonates with previous studies in life history theory [32, 33]. In the language of this field, our fragmentation strategies specify both the size at first reproduction and clutch size, where the latter is subject to a very specific trade-off between the number and size of offspring mathematically given by integer partitions.

We found that for any fitness landscape, the optimal life cycle is always a deterministic fragmentation mode involving the regular schedule of group development and fragmentation. This makes intuitive sense given our assumption that the environment is constant. However, this result might not hold if the environment is variable so that the fitness landscape changes over time. In this case different pure fragmentation modes will be optimal at different times, and natural selection might favour life cycles that randomly express a subset of locally optimal fragmentation patterns. Indeed, the evolution of variable phenotypes in response to changing environmental conditions (also known as bet hedging [34, 35]) has been demonstrated in other life history traits, such as germination time in annual plants [36], and capsulation in bacteria [37]. The extent to which mixed fragmentation modes can evolve via a similar mechanism is beyond the scope of this paper, but it can be addressed in future work by applying existing theory on matrix population models in stochastic environments [22].

We found that when fragmentation is costless, only strategies involving binary splitting (i.e., fragmentation into exactly two parts) are optimal. This result holds for all possible fitness landscapes, and hence any specification of how fecundity or survival benefits might accrue to group living. In particular, the optimal fragmentation mode under monotonic fitness landscapes is generally one of two types: equal binary fragmentation, which involves fission into two equal size groups, and the unicellular propagule strategy, which involves the production of two groups, one comprised of a single cell. Equal fragmentation is favoured when there is a significant advantage associated with formation of even the smallest group, whereas production of a unicellular propagule is favoured when the benefits associated with group size are not evident until groups become large. This makes intuitive sense: when advantages arise when groups are small, it pays for offspring to be in groups (and not single cells). Conversely, when there is little gain until group size is large, it makes sense to maintain one group that reaps this advantage. Interestingly, two bacteria that form groups and are well studied from a clinical perspective, *Neisseria gonorrhoeae* and *Staphylococcus aureus*, both show evidence of the above binary splitting fragmentation modes: *Neisseria gonorrhoeae* divide into groups of two equal sizes [6], while *Staphylococcus aureus* divide into one large group plus a unicellular propagule [7]. This leads to questions concerning the nature of the fitness landscape occupied by these bacteria and the basis of any collective level benefit as assumed by our model.

Once cell loss upon fragmentation is incorporated as a factor in collective reproduction, a wider range of fragmentation patterns becomes optimal. When fragmentation costs are fixed to a given number of cells, optimal fragmentation modes include those where splitting involves the production of multiple offspring. Among these, a prominent fragmentation strategy is multiple fission, where a group breaks into multiple independent cells. Such a fragmentation mode is reminiscent of palintomy in the volvocine algae[38]. A key difference between our “multiple fission” and palintomy is that the former involves a group of cells growing up to a threshold size at which point fragmentation happens, while the latter involves a single reproductive cell growing to many times its initial size and then undergoing several rounds of division. However, reinterpreting birth rates of cells in groups as growth rates of unicells of different sizes allows us to use our analysis to determine conditions under which such a mode of fragmentation is more adaptive than, say, the more standard strategy of growing to twice the initial size and then dividing in two (which for arbitrary sizes of offspring groups is equivalent to our “equal binary fragmentation” mode). Our results suggest that palintomy is favored over binary fission (and any other fragmentation mode) under a wide range of demographic scenarios (Fig. 6C).

Many multicellular organisms are characterised by a life cycle whereby adults develop from a single cell [39]. Passing through such a unicellular bottleneck is a requirement for sexual reproduction based on syngamy, but life cycles with unicellular stages are also common in asexual reproduction modes such as those used by multicellular algae and ciliates [40], and colonial bacteria such as *S. aureus* [7]. If multicellularity evolved because of the benefits associated to group living, why do so many asexual multicellular organisms begin their life cycles as solitary (and potentially vulnerable) cells? Explanatory hypotheses include the purge of deleterious mutations and the reduction of within-organism conflict [41, 39]. Our results make the case for an alternative (and perhaps more parsimonious) explanation: often, a life cycle featuring a unicellular bottleneck is the best way to guarantee that the “parent” group remains as large as possible to reap maximum fecundity and/or survival advantages of group living. Indeed, our theoretical results resonate with previous experimental work demonstrating that single-cell bottlenecks can be adaptive simply because they constitute the life history strategy that maximises reproductive success [42].

Previous theoretical work has explored questions related to the evolution of multicellularity using matrix population models similar to the one proposed in this paper. In a seminal contribution, Roze and Michod [43] explored the evolution of propagule size in the face of deleterious and selfish mutations. In their model, multicellular groups first grow to adult size and then reproduce by splitting into equal size groups, so that fragmentation mode strategies can be indexed by the size of the propagule. In our terminology, this refers to either “multiple fission” or “multiple groups”. An important finding of Roze and Michod [43] is that, even if large groups are advantageous, small propagules can be selected because they are more efficient at eliminating detrimental mutations. We did not study the effects of mutations, but allowed for general fitness landscapes and fragmentation modes, including cases of asymmetric binary division (e.g., the unicellular propagule strategy) neglected by Roze and Michod [43]. Our results indicate that modes of fragmentation involving single cells can lead to growth rate maximisation even when small propagule sizes divide less efficiently or die at a higher rate. In particular, we have shown that if fragmentation is costly, a strategy consisting of a multiple fragmentation mode with a propagule size of one (i.e., the small propagule strategy studied by Roze and Michod [43]) can be adaptive for reasons other than the elimination of deleterious mutations.

Closer to our work, Tarnita et al. [18] investigated the evolution of multicellular life cycles via two alternative routes: “staying together” (ST, whereby offspring cells remain attached to the parent) and “coming together” (CT, whereby cells of different origins aggregate in a group). In particular, they studied the conditions under which a multicellular strategy that produces groups via ST can outperform a solitary strategy whereby cells always separate after division. The way they modelled group formation and analyzed the resulting population dynamics (by means of biological reactions and matrix models) is closely related to our approach. Indeed, their solitary strategy is our binary mode 1+1, while their ST strategy corresponds to a particular kind of binary mixed fragmentation mode. However, the questions we ask are different. Tarnita et al. [18] were concerned with the conditions under which (multicellular) strategies that form groups can invade and replace (unicellular) strategies that remain solitary. Contrastingly, we aimed to understand the optimal fragmentation mode out of the vast space of fragmentation strategies comprising all possible deterministic and probabilistic pathways by which complexes can stay together and split apart. A key finding is that, for any fitness landscape and if the environment is constant, mixed fragmentation modes such as some of the ST strategies considered by Tarnita et al. [18] will be outperformed by at least one pure fragmentation mode.

More recently, Rashidi et al. [20] developed a conceptual framework to study the competition of life cycles that involved five different life cycles defined by fragmentation patterns of the form 1+1+…+1 and an associated genetic control. Their model, which explicitly considers growing cells of different size, showed that depending on the fitness landscape, each of their five life cycles could prevail. By extending the range of life cycles to encompass all possible fragmentation modes (albeit with less detailed attributes), we have shown that certain life cycles will be suboptimal for any given fitness landscape.

In line with many studies in life history theory [32, 33], we made the simplifying assumption that the phenotype consists of demographic traits (in our case, probabilities of fragmenting into given fragmentation patterns) linked by trade-offs which interact to determine fitness (growth rate). This allowed us to predict the optimal phenotype at equilibrium at the expense of leaving unspecified whether, due to genetic constraints, such an equilibrium will be possible in an actual biological system. The question that inevitably arises is whether, given a presumptive genotype-phenotype mapping, it is possible for evolution to fine tune life cycles with group-level properties (such as specific fragmentation patterns) so that optimal fragmentation modes will be obtained as the endpoint of an evolutionary process. While a complete answer requires a more sophisticated analysis, we see no conceptual obstruction preventing seemingly arbitrary fragmentation modes to evolve. Firstly, genotype-phenotype maps of existing organisms can be complex and offer opportunity for adaptation, involving important qualitative behavioral changes [44, 45, 46]. Secondly, small genotypic changes can produce major phenotypic changes. For instance, Hammerschmidt et al. [3] observed the emergence of collective-level properties in a previously unicellular organism that was caused by a small number of mutations. Thirdly, even if a current set of genes cannot provide an appropriate template for given phenotypic traits, new genes can emerge de novo [47, 48, 49, 50, 51]. Finally, theoretical arguments suggest that genetic constraints can be effectively overcome in phenotypic evolution provided there is a rich variety of new mutant alleles [52]. We thus think that, both in the field and in the laboratory, multicellular organisms will be able to evolve a phenotype close to the optimal fragmentation mode in the (very) long run.

## Acknowledgements

We would like to thank two anonymous reviewers for their helpful comments. Appendix

## Appendix

## A Mixed fragmentation modes

A mixed fragmentation mode assigns a probability *q*_*κ*_ to each possible fragmentation pattern (or partition) *κ* ⊦ 2,…, *κ* ⊦ *n*, where *n* is the maximum group size. Such probabilities satisfy Σ_*κ*⊦*j*_*q*_*κ*_ = 1 for *j* = 2,…, *n*, i.e., when growing from size *j* – 1 to *j* one of the partitions *κ* ⊦ *j* (including staying together without splitting, *κ* = *j*) will certainly occur. Additionally, we impose *q*_*n*_ = 0 so that, when growing from size *n* – 1 to size *n*, a group can no longer stay together and will necessarily fragment. It follows that a given life cycle or fragmentation mode can be represented by a set of vectors of the form

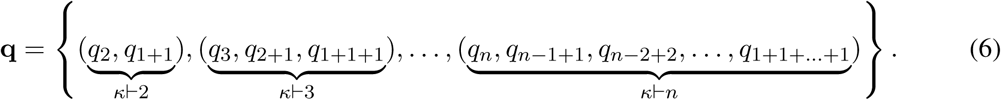

Pure life cycles are a particular case where splitting probabilities *q*_*κ*_ are either zero or one, so that only one fragmentation pattern with more than one offspring group occurs.

A mixed life cycle can be understood as a set of reactions. A number *n* – 1 of reactions, of the type

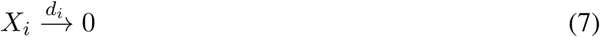

model the death of groups; these are independent of the fragmentation mode. An additional number of reactions, one per each non-zero element of the vector **q**, models the birth of units and the growth or fragmentation of groups. These reactions are of the type

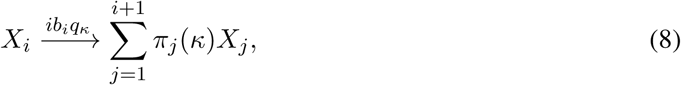

whereby a group of size *i* turns into a group of size *i* + 1 at rate *ib*_*j*_, and then instantly divides with probability *q*_*κ*_ into offspring groups in a way described by fragmentation pattern *κ* ⊦ *i* + 1, where parts equal to *ℓ* appear a number *π_ℓ_*(*κ*) of times. These reactions depend on the life cycle, which specifies the probabilities of fragmentation patterns. For instance, the reaction

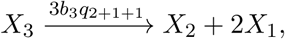

stipulates that groups of size 3, which grow to size 4 at rate 3*b*_3_, will split with probability *q*2+1+1 into one group of size 2 and two groups of size 1. The growth of a group without fragmentation is also), incorporated in the set of reactions given by (8). For instance, the reaction

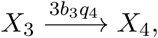

stipulates that groups of size 3, which grow to size 4 at rate 3*b*_3_, will not split with probability *q*_4_.

The sets of reactions (7) and (8) give rise to the system of differential equations

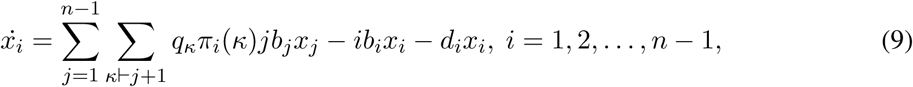

where *x*_*i*_ denotes the abundance of groups of size *i*. This linear system can be represented in matrix form as

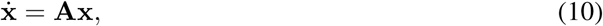

where x = (*x*_1_, *x*_2_,…, *x*_*n*–1_) is the vector of abundances of the groups of different size and **A** is a (*n* – 1) × (*n* – 1) matrix with elements given by

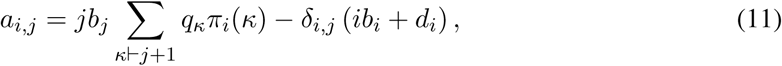

where *δ*_*i*,*j*_ is the Kronecker delta. Since *π*_*i*_(*κ*) = 0 for *κ* ⊦ *j* + 1 and *i* > *j* + 1 (a partition of a number has no parts larger than the number), the entries of **A** below the subdiagonal are zero. As an example, consider *n* = 4. The projection matrix for this case is given by

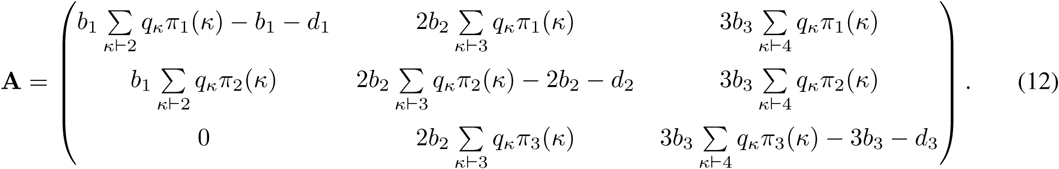

## B Mixed fragmentation modes are dominated

For any fitness landscapes, mixed fragmentation modes are dominated by at least one pure life cycle. In other words, the optimal life cycle is pure.

To prove this result, consider the set of partitions *κ* ⊦ *j* for a given *j*, fix the probabilities of fragmentation patterns *ν* ⊦ *i* ≠ *j* to arbitrary values, and focus attention on the function

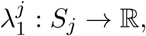

mapping probability distributions in the *ζ*_*j*_-simplex *S*_*j*_ ⊂ ℝ^*ζ*_*j*_^ (specifying the probabilities of all partitions *κ* ⊦ *j*) to the dominant eigenvalue 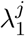 of the associated projection matrix **A**. Our goal is to show that, for any *j*, 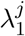 is a quasiconvex function, i.e., that

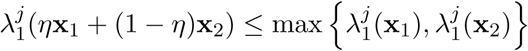

holds for all x_1_, x_2_ ∈ *S*_*j*_ and *η* ∈ [0,1]. Quasiconvexity of 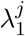 implies that 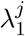 achieves its maximum at an extreme point of *S*_*j*_, i.e., at a probability distribution that puts all of its mass in a single fragmentation pattern. Quasiconvexity of 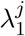 for all *j* then implies that the maximum growth rate λ_1_ is achieved by a pure fragmentation mode, and that mixed fragmentation modes are dominated.

To show that 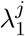 is quasiconvex, we restrict the function to an arbitrary line and check quasiconvexity of the resulting scalar function [53, p. 99]. More precisely, we aim to show that the function

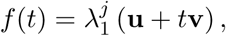

is quasiconvex in *t* for any **u** ∈ *S*_*j*_ and **v** ∈ ℝ^*ζ*_*j*_^ such that **u** + *t***v** ∈ *S*_*j*_. We hence need to verify that

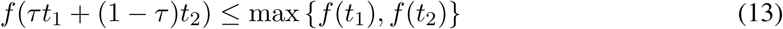

holds for *τ* ∈ [0,1].

To show this, note that the function 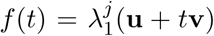 is given implicitly by the largest root of the characteristic polynomial

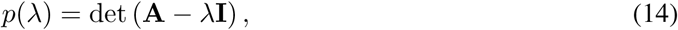

where the probabilities of fragmentation specified by **u** + *t***v** appear in the (*j* – 1)-th column of the projection matrix A (see Eqs. (11) and (12)).

The right hand side of Eq. (14) can be written using a Laplace expansion along the (*j* – 1)-th column of **A** – λ**I**:

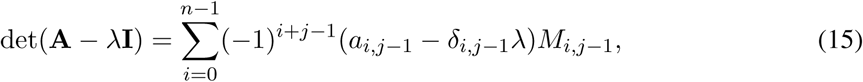

where *δ*_*i*, *j*–1_ is the Kronecker delta and *M*_*i*,*j* – 1_ is the (*i*, *j* – 1) minor of **A**, i.e., the determinant of the submatrix obtained from **A** by deleting the *i*-th row and (*j* – 1)-th column. Each minor *M*_*i*,*j* – 1_ is independent of *t* because the only entries of **A** that depend on *t* appear in the (*j* – 1)-th column. Moreover, each entry *a*_*i*, *j*–1_ is either zero or a linear function of t. Hence, *p*(λ) is a polynomial on λ with coefficients that are linear in *t*, i.e., of the form

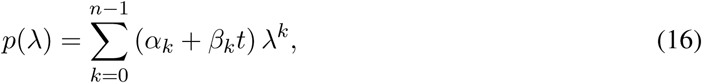

for some *α*_*k*_, *β*_*k*_. Moreover, since the leading coefficient must be (—1)^*n*–1^ (the matrix **A** is (*n* – 1) × (*n* – 1)), it follows that *α*_*n*–1_ = (–1)^*n*–1^ and *β*_*n*–1_ = 0.

Denote by *p*_*τ*_(λ), *p*_1_(λ), and *p*_2_(λ) the characteristic polynomials corresponding to, respectively, the probability distributions given by **u** + [*τt*_1_ + (1 – *τ*)*t*_2_] **v**, **u** + *t*_1_ **v**, and **u** + *t*_2_ **v**. From Eq. (16), these are given by

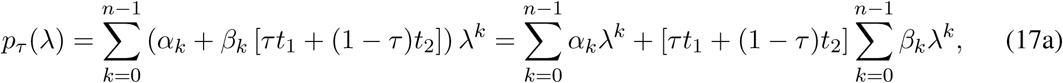

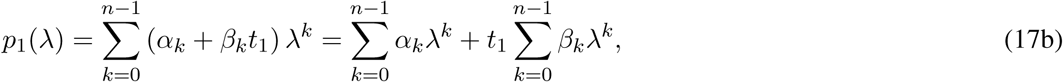

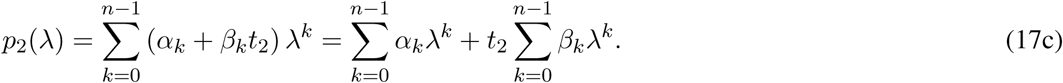

Subtracting Eq. (17b) from Eq. (17a), and Eq. (17c) from Eq. (17a), we can write

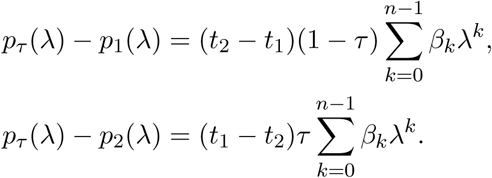

Note that the signs of these differences are always different, i.e., either (i) *p_τ_*(λ) – *p*_1_(λ) ≥ 0 and *p_τ_*(λ) – *p*_2_(λ) ≤ 0, or (ii) *p_τ_*(λ) – *p*_1_(λ) ≤ 0 and *p_τ_*(λ) – *p*_2_(λ) ≥ 0. In the first case, we have *p*_1_(λ) ≤ *p_τ_*(λ) ≤ *p*_2_(λ) and in the second we have *p*_2_(λ) ≤ *p_τ_*(λ) ≤ *p*_1_(λ), i.e., for each λ, *p_τ_*(λ) lies between *p*_1_(λ) and *p*_2_(λ), or, equivalently

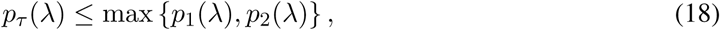

for all λ. Since 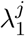 is the largest root of *p*(λ), and since *p_τ_*(λ), *p*_1_ (λ), and *p*_2_(λ) all have the same sign in the limit when λ tends to infinity (their leading coefficients are all equal to *α*_*n*–1_ = (—1)^*n*–1^), condition (18) implies condition (13), thus proving our claim. See Fig. 8 for an illustration.

**Figure 8:**
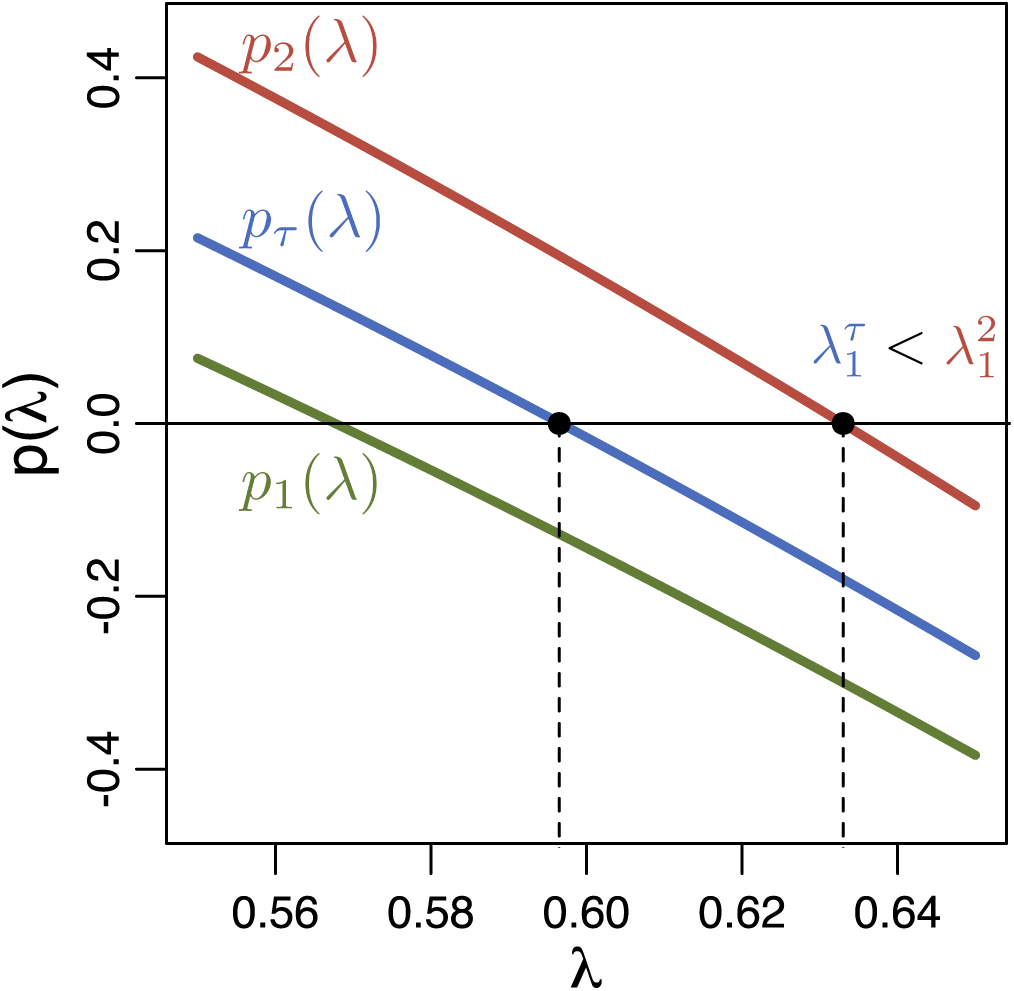
**Population growth rate** λ_1_ **is quasiconvex.** Consider two fragmentation modes *q*_1_ and *q*_2_ which differ only in the probabilities of fragmentation patterns at a single size *j*. Then, for any 0 ≤ *τ* ≤ 1 and corresponding fragmentation mode **q**_*τ*_ = *τ***q**_1_ + (1 – *τ*)**q**_2_, the polynomials *p*(λ) given by Eq. (14) satisfy either *p*_1_(λ) ≤ *p_τ_*(λ) ≤ *p*_2_(λ) or *p*_2_(λ) ≤ *p_τ_*(λ) ≤ *p*_1_(λ). Thus, **q**_*τ*_ leads to a lower growth rate than either **q**_1_ or **q**_2_, i.e., either 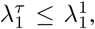 or 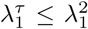 holds. Here, *j* = 3, **q**_1_ = {(0.9,0.1), (<u>0.5,0.5,0</u>), (0,0,0,1,0)}, **q**_2_ = {(0.9,0.1), (<u>0.5,0,0.5</u>), (0,0,0,1,0)}, and *τ* = 0.6. The fitness landscape is given by *b*_*i*_ = 1/*i*, *d*_*i*_ = 0 for all *i*.

## C Mixing between 1+1 and 2+1 is dominated

To show that the life cycle mixing between fragmentation modes 1+1 and 2+1 with probability *q* represented in vector form as

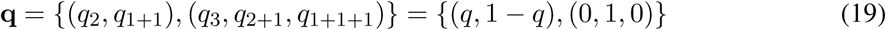

is dominated, consider its growth rate 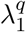 as a function of *q*, as given by the solution of characteristic equation

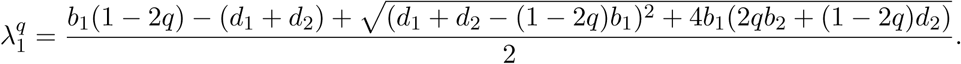

We have 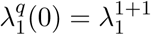 and 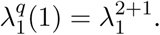 A sufficient condition for *q* to be dominated by either 1+1 or 2+1 is then that 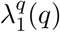 is monotonic in *q*. To show that this is the case, note that the derivative of 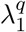 with respect to *q* is given by

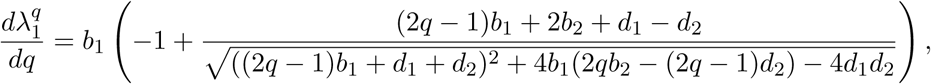

and that such expression is equal to zero if and only if

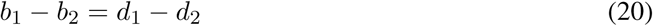

which is independent of *q*. It follows that 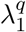 is either nonincreasing or nondecreasing in *q*, and hence that it attains its maximum at either *q* = 0, *q* = 1, or (when (20) is satisfied) at any *q* ∈ [0,1]. Hence, *q* is dominated by either 1+1 or 2+1.

## D Characteristic equation of a pure fragmentation mode

Consider the pure fragmentation mode *κ* ⊦ *ℓ*, whereby groups grow up to size *ℓ* and then fragment according to fragmentation pattern *κ*. The projection matrix is a (*ℓ* – 1) × (*ℓ* – 1) matrix of the form

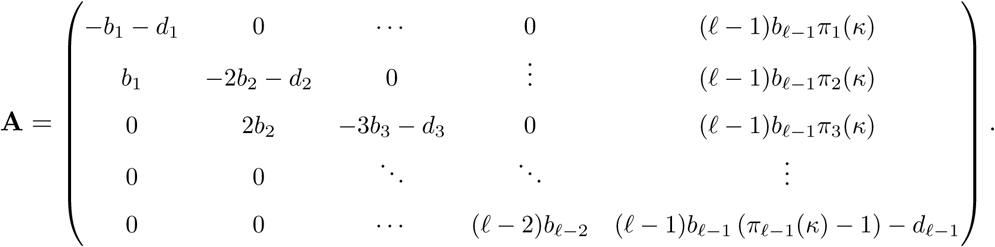

The population growth rate is given by the leading eigenvalue λ_1_ of **A**, i.e., the largest solution of the characteristic equation

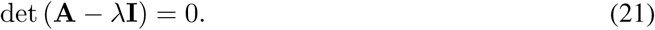

By using a Laplace expansion along the last column of **A** – λ**I**, we can rewrite the left hand side of the above expression (i.e., the characteristic polynomial of **A**) as

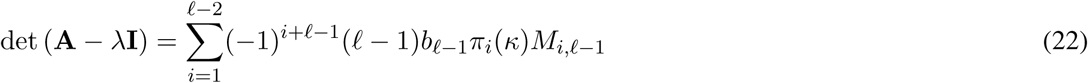

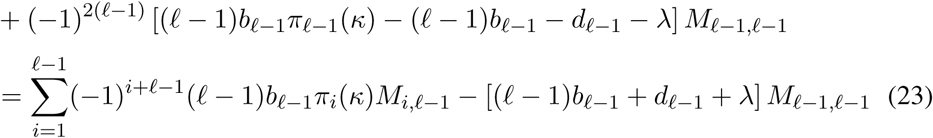

where *M*_*i*, *ℓ* – 1_ is the (*i*, *ℓ* – 1)-th minor of **A** – λ**I**. For all *i* = 1,…, *ℓ* – 1, the minor *M*_*i*, *ℓ* – 1_ is the determinant of a block diagonal matrix, and hence equal to the product of the determinants of the diagonal blocks. Moreover, each diagonal block is either a lower triangular or an upper triangular matrix, whose determinant is given by the product of the elements in their main diagonals. We can then write

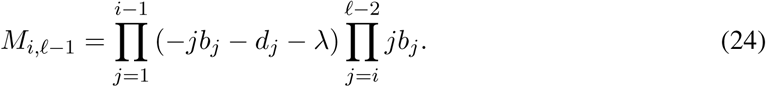

Substituting Eq. (24) into Eq. (23) and simplifying, we obtain

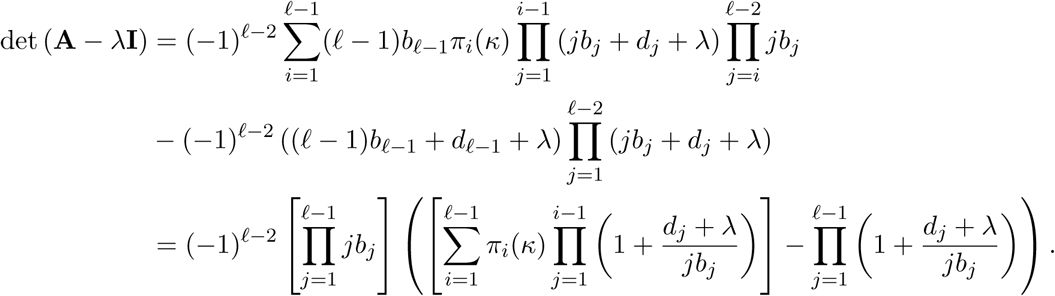

Replacing this expression into the characteristic equation (21), dividing both sides by 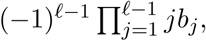 and simplifying, we finally obtain that the characteristic equation (21) can be written as

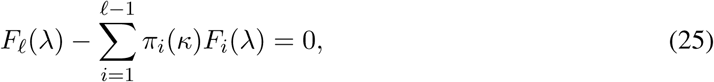

where

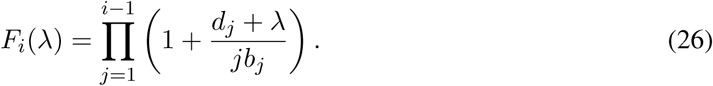

Note that the following two transformations:

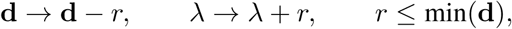

and

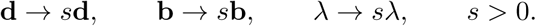

preserve the solution of Eq. (25) This allows us to set *b*_1_ = 1 and min(**d**) = 0 without loss of generality.

## E Fragmentation modes are dominated by binary splitting

We can show that, for any fitness landscape, binary fragmentation achieves a larger growth rate than splitting into more than two offspring groups. To prove this, consider (i) positive integers *m*, *j*, and *k* such that *m* > *j*+*k*, (ii) an arbitrary partition *τ* ⊦ *m*–*j*–*k*, and (iii) the following three fragmentation modes:

1. *κ*_1_ = *j* + *k* + *τ* ⊦ m, whereby a complex of size m fragments into one complex of size *j*, one complex of size *k*, and a number of offspring complexes given by partition *τ*.
2. *κ*_2_ = (*j* + *k*) + *τ* ⊦ *m*, whereby a complex of size *m* fragments into one complex of size *j* + *k*, and a number of offspring complexes given by partition *τ*.
3. *κ*_3_ = *j* + *κ* ⊦ (*j* + *k*), a binary splitting fragmentation mode whereby a complex of size *j* + *k* fragments into two offspring complexes: one of size *j*, and one of size *k*.

Fragmentation mode *κ*_1_ leads to a number of offspring groups equal to

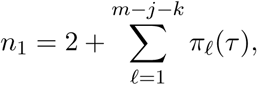

fragmentation mode *κ*_2_ to a number of offspring groups equal to

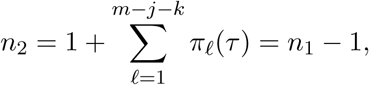

and fragmentation mode *κ*_3_ to a number of offspring groups equal to two. Denoting by 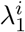 the growth rate of fragmentation mode *κ*_*i*_, we can show that, for any fitness landscape, either 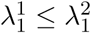 or 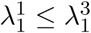 holds, i.e., a fragmentation mode with more than two parts is dominated by either a fragmentation mode with one part less or by a fragmentation mode with exactly two parts. By induction, this implies that the optimal life cycle is always one within the class of binary fragmentation modes.

To prove that either 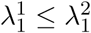 or 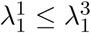 holds, let us denote by *p*_*i*_(λ) the characteristic polynomial associated to mode *κ*_*i*_, as given by the left hand side of Eq. (25) after the replacement *κ* = *κ*_*i*_. The growth rate 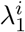 of mode *κ*_*i*_ is hence the largest root of *p*_*i*_(λ). The polynomials *p*_1_(λ), *p*_2_(λ), and *p*_3_(λ) are then given by

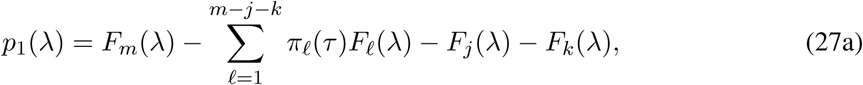

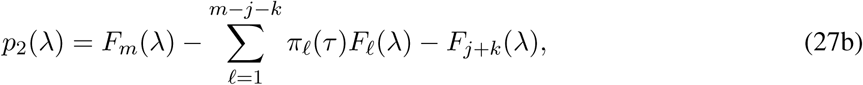

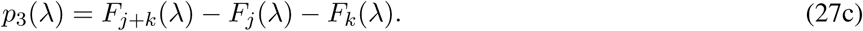

These polynomials satisfy the following two properties. First,

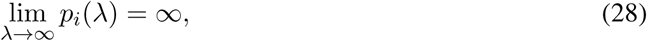

as the leading coefficient of the left hand side of Eq. (25) is always positive. Second, we can write

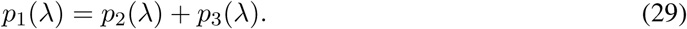

Now, evaluating Eq. (29) at 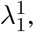 and since 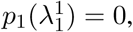 it follows that 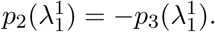 Hence, only one of the following three scenarios is satisfied: (i) 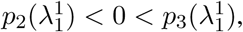 (ii) 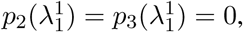 or (iii) 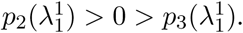 If 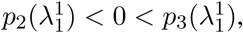 and by Eq. (28) and Bolzano’s theorem, 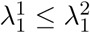 holds. Likewise, if 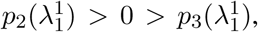 then 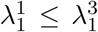 holds. Finally, if 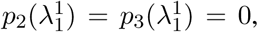 then both 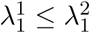 and 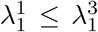 hold. See Fig. 9 for a graphical illustration of these arguments. We conclude that either 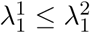 or 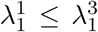 must hold, which proves our result.

**Figure 9:**
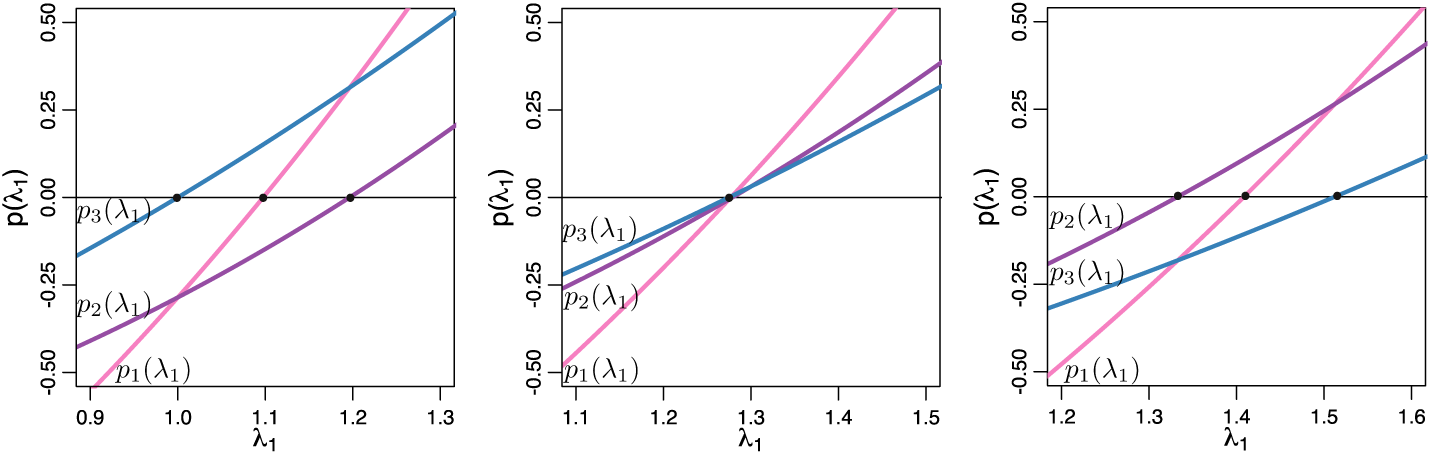
**The population growth rate induced by a fragmentation mode with more than two offspring groups is dominated.** Consider the characteristic polynomials *p*_*i*_(λ_1_) for partitions *κ*_1_ = 2 + 1 + 1, *κ*_2_ = 3 + 1 and *κ*_3_ = 2 + 1. *Left:* Fitness landscape **b** = (1,1,1.4), **d** = (0,0,0). Since 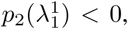 *κ*_1_ is dominated by 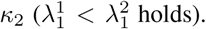 *Center.* Fitness landscape 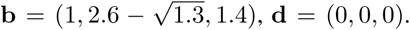 Since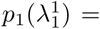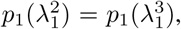 *κ*_1_ is weakly dominated by 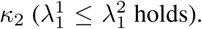 *Right*: Fitness landscape **b** = (1,1.9,1.4), d = (0,0,0). Since 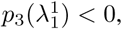 *κ*_1_ is dominated by 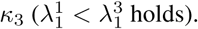

## F Optimality maps for *n* = 4

For *n* = 4 there are four pure fragmentation modes: 1+1, 2+1, 2+2, and 3+1. From Eq. (25), their characteristic polynomials are respectively given by

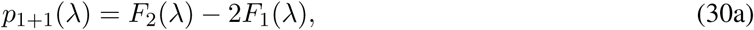

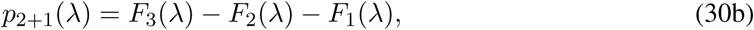

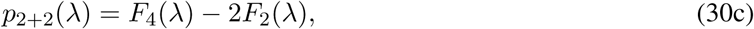

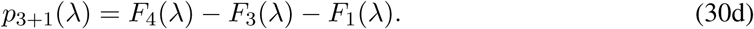

The optimality maps shown in Fig. 3 of the main text were obtained by comparing the largest root of these characteristic polynomials, which we computed numerically. For fecundity landscapes, we tested fitness landscapes of the form {**b**, **d**} = {(1, *b*_2_, *b*_3_), (0,0,0)} with *b*_2_ and *b*_3_ taken from a rectangular grid of size 300 by 300 with *b*_2_ ∈ [0, 5] and *b*_3_ ∈ [0, 5]. For viability landscapes, we tested fitness landscapes of the form {**b**, **d**} = {(1,1,1), (5, *d*_2_, *d*_3_)} with *d*_2_ and *d*_3_ taken from a rectangular grid of size 300 by 300 with *d*_2_ ∈ [0,10] and *d*_3_ ∈ [0,10].

The boundaries between areas of optimality can still be computed analytically. They are given by the fitness landscapes at which two fragmentation modes have the same population growth rate.

The following are the boundaries between areas of optimality under fecundity fitness landscape (assuming *b*_1_ = 1 for simplicity):

- Between fragmentation modes 1+1 and 2+1: *b*_2_ = 1, *b*_3_ ≤ 1.
- Between fragmentation modes 1+1 and 3+1: 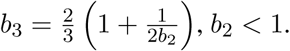
- Between fragmentation modes 2+1 and 2+2: 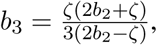 where 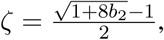 and *b*_2_ > 1.
- Between fragmentation modes 3+1 and 2+2: 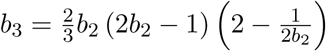 and *b*_2_ > 1

The following are the boundaries between areas of optimality under viability fitness landscape (assuming *d*_1_ = 0 for simplicity):

- Between fragmentation modes 1+1 and 2+1: *d*_2_ = 0, *d*_3_ > 0.
- Between fragmentation modes 1+1 and 3+1: 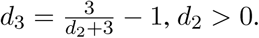
- Between fragmentation modes 2+1 and 2+2: 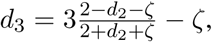 where 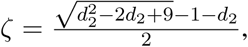 and *d*_2_ < 0.
- Between fragmentation modes 3+1 and 2+2: 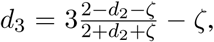 where 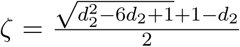 and *d*_2_ < 0

## G Costly fragmentation

For costly fragmentation, some cells are lost upon the fragmentation event. In this case the biological reactions are still given by Eqs. (7) and (8). However, under costly fragmentation the sum of sizes of offspring groups is smaller than the size of the parent group. Therefore, in Eq. (8), *κ* is a partition of *i'* ≤ *i* + 1 (and not strictly of *i* + 1 as it was under costless fragmentation). Indeed, *i'* = *i* + 1 in the case of trivial partitions with one part (when a group grows without splitting), but *i'* < *i* + 1 for nontrivial partitions with two or more parts (where the group grows in size by one cell and then splits). In this latter case, *i'* = *i* – *π* + 2 (where *π* is the number of offspring groups) for the case of proportional costs, and *i'* = *i* for the case of fixed costs.

To illustrate the difference in the available sets of partitions for each of the three scenarios we investigate (costless fragmentation, fragmentation with proportional cost, fragmentation with fixed cost), consider the following possible reactions for a 4-cell group growing into a 5-cell group. For costless fragmentation, we have

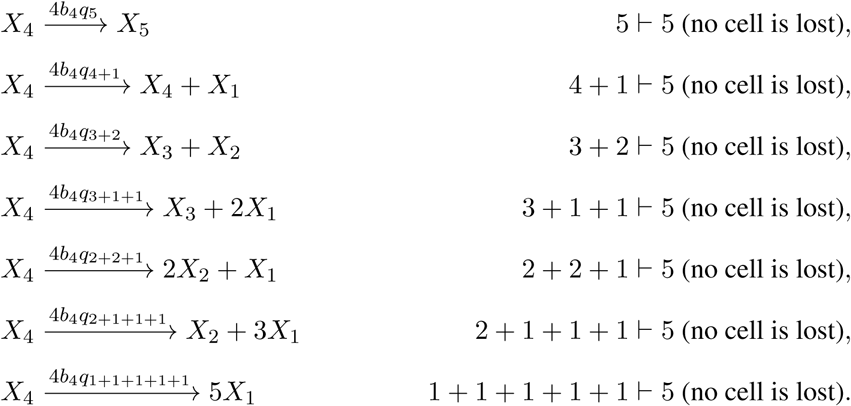

For fragmentation with fixed cost, we have

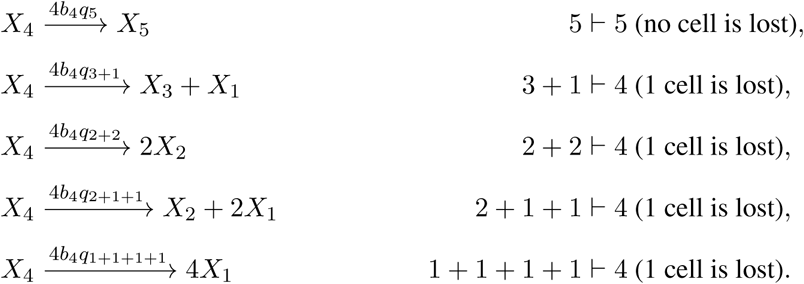

Finally, for fragmentation with proportional cost, we have

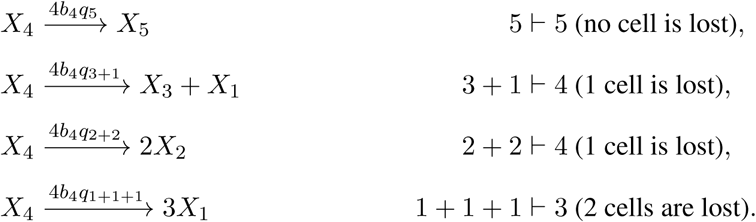

The combined probability of all outcomes of aggregate growth must be equal to one. In the case of costless fragmentation, this condition has been given by Σ_*κ*⊦*i*+1_ *q*_*κ*_ = 1 for *i* = 1,…, *n* – 1. For costly fragmentation this condition changes to Σ_*κ*⊦*i'*_ *q*_*κ*_ = 1 for *i* = 1,…, *n* – 1, with *i'* as defined above. The expressions for the system of differential equations and the projection matrix for general mixed strategies (Eqs. (9) and (12)) are changed accordingly. For pure fragmentation modes, the projection matrix given in the main text and the characteristic equation given in Eq. (25) remain valid, but *κ* is no longer a partition of *i* + 1 but of *i'* as defined above.

## H With proportional costs, fragmentation modes are dominated by binary splitting

For fragmentation with proportional costs, a group fragmenting into *π* offspring groups incurs a cost of *π* – 1 cells. In this case, similarly to the case for costless fragmentation, nonbinary fragmentation modes are dominated by binary fragmentation modes. To prove this, consider (i) positive integers *m*, *j*, and *k* such that *m* > *j* + *k* + 4, (ii) an arbitrary partition *τ* with *π* ≥ 2 parts such that *τ* ⊦ *m* – *j* – *k* – *π* – 2, and (iii) the following three fragmentation modes:

1. *κ*_1_ = *j* + *k* + *τ* ⊦ *m* – *n* – 1, whereby a complex of size *m* fragments into one complex of size *j*, one complex of size *k*, and *π* complexes given by partition *τ*, and *π* + 1 cells die.
2. *κ*_2_ = (*j* + *k* + 1) + *τ* ⊦ *m* – *π*, whereby a complex of size *m* fragments into one complex of size *j* + *k* + 1 and *π* complexes given by partition *τ*, and *π* cells die.
3. *κ*_3_ = *j* + *κ* ⊦ (*j* + *k*), a binary fragmentation mode whereby a complex of size *j* + *k* + 1 fragments into two offspring complexes (one of size *j* and one of size *k*), and one cell dies.

Note that fragmentation mode *κ*_1_ leads to *π* + 2 offspring groups, fragmentation mode *κ*_2_ leads to *π* + 1 offspring groups, and fragmentation mode *κ*_3_ leads to a number of offspring groups equal to two. The rest of the proof is analogous to the one given in Appendix E for the case of costless fragmentation and will be omitted.

